# Collagen IV of basement membranes: I. Origin and diversification of COL4A genes enabling animal evolution and adaptation

**DOI:** 10.1101/2023.10.18.563013

**Authors:** Patrick S. Page-McCaw, Elena N. Pokidysheva, Carl E. Darris, Sergei Chetyrkin, Aaron L. Fidler, Julianna Gallup, Aspirnaut Team, Prayag Murawala, Julie K. Hudson, Sergei P. Boudko, Billy G. Hudson

## Abstract

Collagen IV is a major component of basement membranes, a specialized form of extracellular matrix that enabled the assembly of multicellular epithelial tissues. In mammals, collagen IV assembles from a family of six α-chains (α1 to α6), forming three supramolecular scaffolds: Col-IV**^α121^**, Col-IV**^α345^** and Col-IV**^α121-α556^**. The α-chains are encoded by six genes (COL4A1-6) that occur in pairs in a head-to-head arrangement. In Alport syndrome, variants in COL4A3, 4 or 5 genes, encoding Col-IV**^α345^** scaffold in glomerular basement membrane (GBM), the kidney ultrafilter, cause progressive renal failure in millions of people worldwide. How variants cause dysfunction remains obscure. Here, we gained insights into Col-IV**^α345^** function by determining its evolutionary lineage, as revealed from phylogenetic analyses and tissue expression of COL4 gene-pairs. We found that the COL4A⟨1|2⟩ gene-pair emerged in basal Ctenophores and Cnidaria phyla and is highly conserved across metazoans. The COL4A⟨1|2⟩ duplicated and arose as the progenitor to the COL4A⟨3|4⟩ gene-pair in cyclostomes, coinciding with emergence of kidney GBM, and expressed and conserved in jawed-vertebrates, except for amphibians, and a second duplication as the progenitor to the COL4A⟨5|6⟩ gene-pair and conserved in jawed-vertebrates. These findings revealed that Col-IV**^α121^** is the progenitor scaffold, expressed ubiquitously in metazoan basement membranes, and which evolved into vertebrate Col-IV**^α345^** and expressed in GBM. The Col-IV**^α345^** scaffold, in comparison, has an increased number of cysteine residues, varying in number with osmolarity of the environment. Cysteines mediate disulfide crosslinks between protomers, an adaptation enabling a compact GBM that withstands the high hydrostatic pressure associated with glomerular ultrafiltration.

## Introduction

Collagen IV (Col-IV) is a principal component of basement membranes, a specialized form of extracellular matrix that enabled the genesis and evolution of multi-cellular epithelial tissues (1,2). In pioneering studies of the glomerular basement membrane (GBM) of canine and bovine kidneys collagen IV was identified as a novel collagen and shown to be structurally altered in diabetes nephritis DN (3–9). It was first characterized as a supramolecular network of triple helical protomers composed of α1 and α2 chains (10,11). In subsequent studies of the GBM in Goodpasture’s disease and Alport syndrome (Fig. 1A), four additional chains were discovered, α3-α6 (Fig. 1B) (12–20). The chains are encoded by six genes (COL4A1 to COL4A6), which are located in gene-pairs (COL4A⟨1|2⟩, COL4A⟨3/4⟩ and COL4A⟨5/6⟩) in a head-to-head arrangement on three different chromosomes (Fig. 1C) (21,22). In mammals, the α-chains co-assemble into heterotrimers, called protomers, of three distinct molecular compositions: α121, α345 and α565, which in turn assemble into three distinct supramolecular scaffolds, referred to as Col-IV**^121^**, Col-IV**^α34^**^5^, and Col-IV**^α556—α12^**^1^ (23,24).

**Figure 1.**
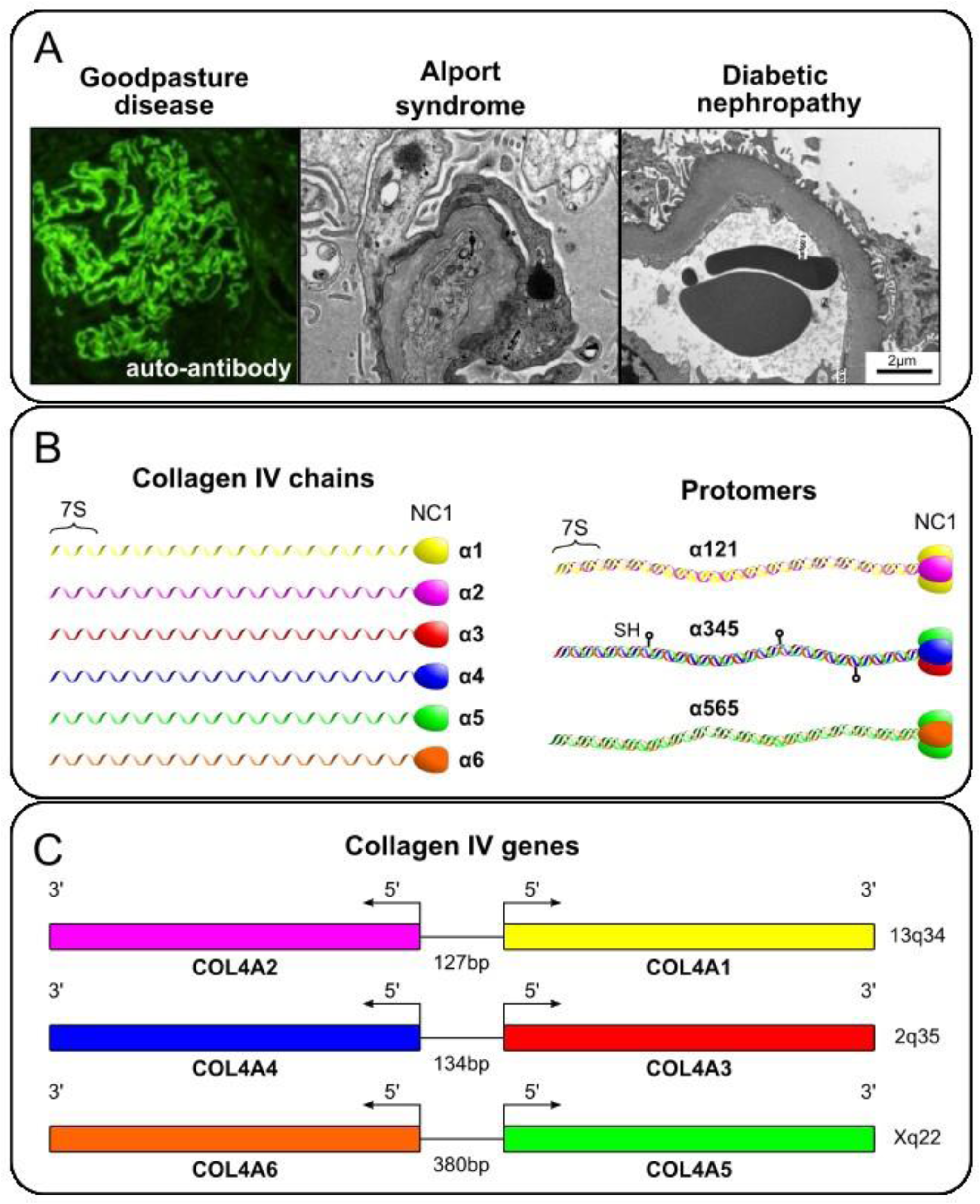
Collagen IV related kidney diseases, protomer and gene organizations in mammals. A. represents characteristic images of GBM abnormalities in three major diseases: Goodpasture disease, an autoimmune disorder, which is diagnosed based on the linear immunofluorescent GBM staining; Alport syndrome, a genetic disorder, where GBM is split with a characteristic basket weaving; Diabetic nephropathy, a common complication of diabetes, where GBM is dramatically thickened. In each of these pathologies collagen IV is affected. B. Six protein chains of collagen IV are produced in mammals. These six chains assemble into trimeric complexes referred to as protomers. The N- and C-terminal 7S and NC1 domains mediate end-to-end homomeric interactions between protomers generating the different Col-IV scaffolds. Both 7S-7S interprotomer tetramers and NC1-NC1 interprotomer dimers are covalently crosslinked. The Col-IV**^a345^** protomer encodes multiple Cys residues which form lateral crosslinks between different Col-IV**^α345^** collagen domains. C. The head-to-head arrangement of the three COL4A gene pairs found in mammals is shown. Their human chromosomal location is indicated and the distances between their transcriptional start sites is shown.

The Col-IV**^α121^** scaffold is a ubiquitous component of mammalian basement membranes. For example, in the nephron this scaffold occurs in the GBM, mesangial matrix, and basement membranes of Bowman’s capsule, tubules and capillaries (Fig. 2) (25,26). The scaffold confers tensile strength and acts as a tether for diverse macromolecules, including laminin, nidogen, proteoglycans, and growth factors, forming the supramolecular complexes that interacts with cell surface receptors (2,27–29). Disrupting this scaffold causes basement membrane destabilization and tissue dysfunction in early mouse development (30). Genetic defects of the Col-IV**^α121^** scaffold cause Gould syndrome and HANAC in humans (31–33).

**Figure 2.**
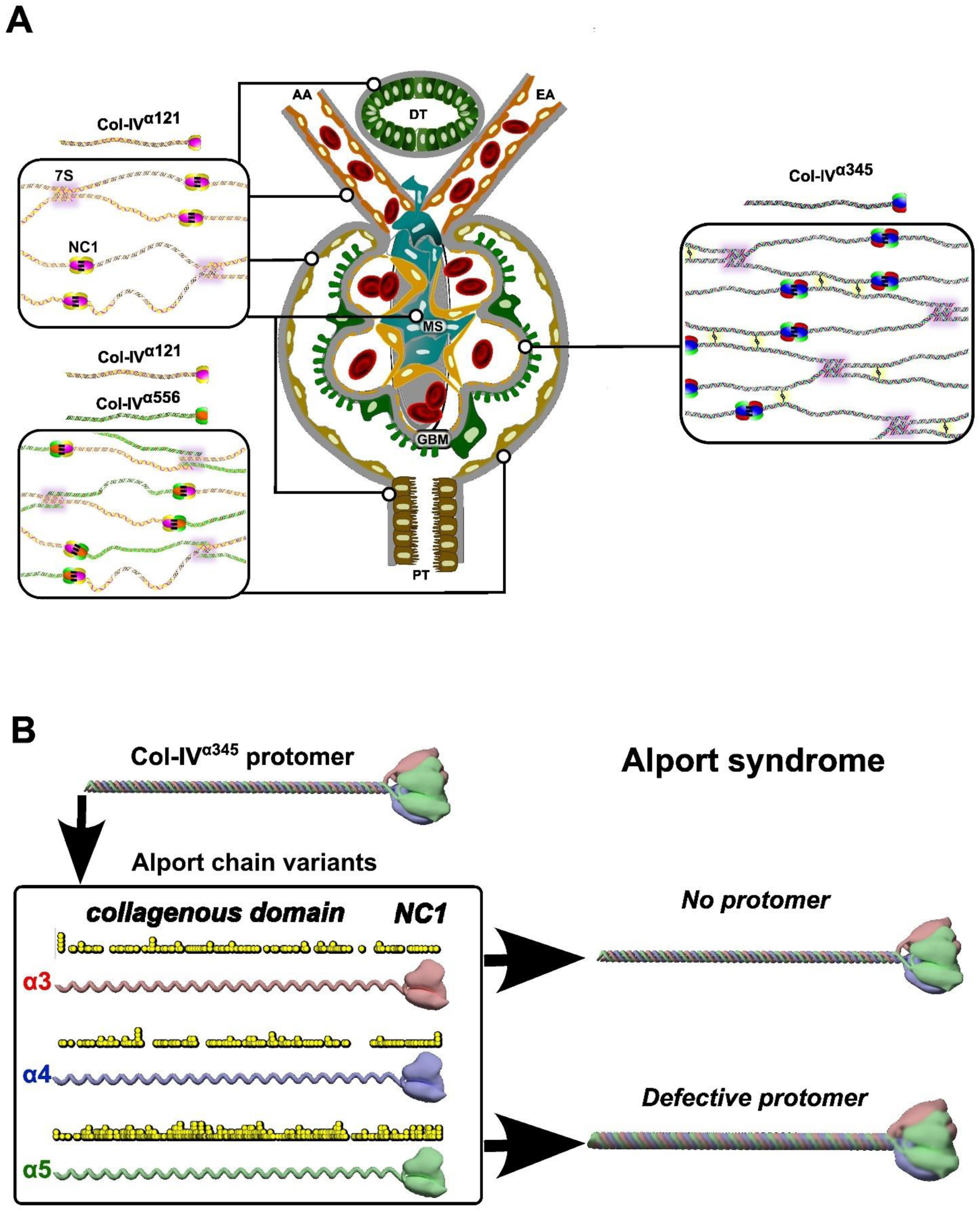
Collagen IV scaffolds of the mammalian kidney glomerulus. A. Three distinct supramolecular Col-IV scaffolds, noted as Col-IV^α**121**^, Col-IV ^α**345**^, and Col-IV ^α**556-** α**121**^, comprise the mammalian kidney glomerulus. Scaffold are assembled from three different triple-helical protomers having three molecular compositions of a-chains: α121, α345 and α565. Protomers are characterized by a 7S domain at the N-terminal, a long collagenous domain of Gly-Xaa-Yaa (GXY) repeats of ∼1400 residues with interruptions in the GXY repeats, followed by a non-collagenous (NC1) domain at the C-terminus of approximately ∼230 residues. B. The Col-IV ^α**121**^ scaffold is a component of the basement membranes surrounding tubules, arterioles and Bowman’s capsule. It is also found within the mesangial space. Uniquely, Bowman’s capsule contains the hetero-scaffold of Col-IV ^α**556-** α**121**^. The glomerular corpuscle consists of looped capillaries lined with fenestrated endothelial cells, the glomerular basement membrane (GBM) and podocytes. The Col-IV^α**345**^ scaffold is the major component of GBM; it is reinforced by disulfide bonds which form lateral crosslinks between protomers. **DT** – distal tubule; **PT** – proximal tubule; **MS** – mesangial space; **AA** – afferent arteriole; **EA** – efferent arteriole; **GBM** – glomerular basement membrane. C. Mutations in any of the COL4A3, A4 or A5 genes that encode the Col-IV ^α**345**^ scaffold cause Alport syndrome. Known disease variants are mapped onto the predicted protein structures as yellow circles. Disease variants cause either the assembly of defective protomers or their complete absence of assembled protomers.

Conversely, the collagen IV**^α345^** scaffold, encoded by COL4A⟨3/4⟩ and COL4A⟨5/6⟩ gene-pairs, has a restricted tissue distribution in the inner ear, testis and kidney (34). In the nephron, collagen IV**^α345^** is the major component of the GBM, a critical morphological feature that functions as an ultrafilter of proteins (Fig. 2) (25,26,34). In Goodpasture’s disease (GP), affecting thousands of people worldwide, autoantibodies target neoepitopes in the Col-IV**^α345^** scaffold causing rapidly progressive renal failure (19,35–41). In diabetic nephropathy (DN), affecting millions of people, this scaffold is also involved in the thickening of the GBM morphology that is associated with progressive kidney failure. In Alport syndrome (AS), affecting millions of people, nearly two thousand genetic variants occur in the COL4A3, COL4A4, and COL4A5 genes(15,42–46). Variants cause either loss of scaffold from the GBM or assembly of a defective scaffold, causing proteinuria and progression to kidney failure (26,47). A knowledge of how the Col-IV**^α345^** scaffold functions at the molecular level is critical to understanding the pathogenesis of GP, DN and AS, thus providing a framework for the development of therapies.

Here, we sought to gain insights into Col-IV**^α345^** function by determining its evolutionary lineage with the Col-IV**^α121^** scaffold, as evinced from phylogenetic analyses, gene synteny and tissue expression of cognate COL4 gene-pairs. The recent availability of a plethora of high-quality genome assemblies provided an approach to trace the evolutionary emergence of COL4 gene-pairs. The findings revealed that Col-IV**^α121^** scaffold is the progenitor, expressed ubiquitously in basement membranes in all animals, and that evolved into vertebrate Col-IV**^α345^** scaffold with expression mainly in the GBM. The Col-IV**^α345^** scaffold differs from Col-IV**^α121^** by an increased number of lateral disulfide crosslinks, indicating an evolutionary adaptation that enabled the assembly of a compact GBM that withstands the high hydrostatic pressure associated with glomerular ultrafiltration.

## Results

### Phylogenetic analysis of the evolutionary lineage of COL4 gene-pairs

We sought to determine the evolutionary lineage of the COL4 gene-pairs, using the approach of comparative genomics and syntenic relationships. DNA sequence is conserved within orthologous and paralogous genes which allows evolutionary relationships between genes to be determined. Analogously, gene content of chromosomes, called synteny, is also conserved even across large evolutionary distances (48,49). Moreover, gene order and orientation, collinearity, is often conserved in microsyntenic blocks. Microsyntenic conservation can provide information about the origin of gene families following speciation and chromosomal duplication. Importantly, recent advances in sequencing technology have facilitated the analysis of gene order and locus structure in diverse species (50). These advances have been accompanied by improvements and standardization in the presentation of genome structure that facilitates an informatics approach to decipher the arrangement and genetic lineage of the COL4 gene family.

Prior studies (21,50,51) established that the six mammalian COL4 genes are arranged in head-to-head pairs, noted as COL4A⟨1|2⟩, COL4A⟨3|4⟩ and COL4A⟨5|6⟩ gene-pairs (Fig. 1C). We searched diverse genomes across metazoa for each of these gene-pairs. First, we found the COL4A⟨1|2⟩ gene-pair to be conserved across metazoans, with only a few exceptions in the Protosome lineage, (Fig.3 and Fig.4). In the nematode clade, for example, the COL4A1 and COL4A2 genes (in *C. elegans* the genes emb-9 and let-2) occur as single, unlinked loci. In other examples, the planarians have multiple copies of single, unlinked COL4 genes, and *Owenia fusiformis* has both the COL4A⟨1|2⟩ gene-pair and a COL4 single gene. Other Protostomes, for example *Schistosoma mansoni*, have gene arrangements similar to that in Ctenophores in which the COL4 loci include a total of four COL4 genes with each set arranged head-to-tail and a COL4A⟨1|2⟩ gene pair positioned adjacent to a single COL4 gene. Second, we found that the COL4A⟨3|4⟩ gene-pair emerged in hagfish and lamprey lineages and was conserved in all vertebrates, except for amphibians. Third, the COL4A⟨5|6⟩ gene-pair emerged in cartilaginous fish and is conserved in all vertebrates (Fig.3). Collectively, our findings indicate that the COL4A⟨1|2⟩ gene-pair is the progenitor of vertebrate COL4A⟨3|4⟩ and that either COL4A⟨1|2⟩ or COL4A⟨3|4⟩ is the progenitor of the COL4A⟨5|6⟩ gene-pair.

**Figure 3.**
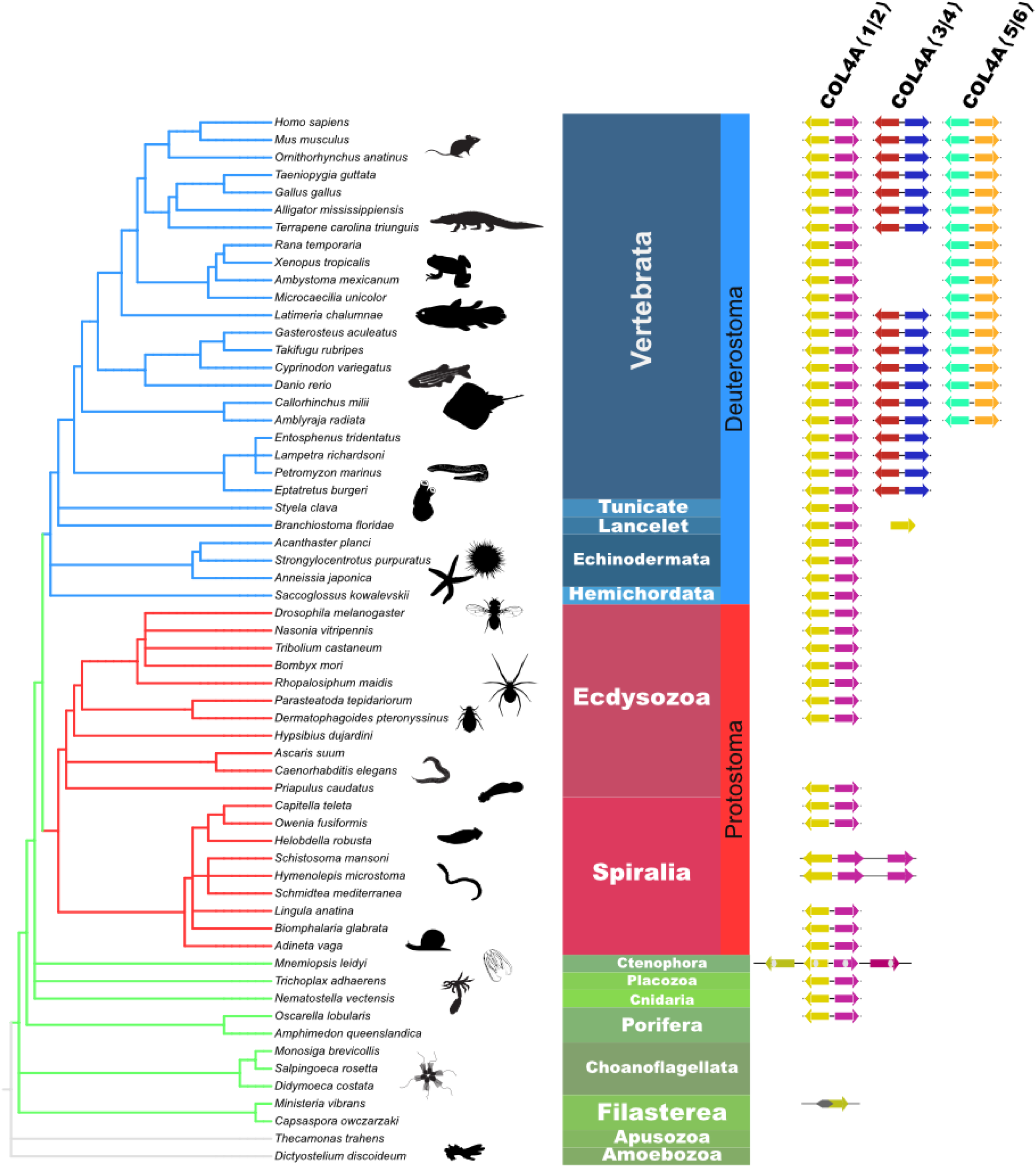
An evolutionary lineage of COL4A genes that gave rise to the appearance of the COL4A (3|4) and COL4A (5|6) gene pairs by gene duplication in vertebrate animals. The eukaryotic cladogram with selected species showing evolution of the COL4A gene pair family mapped onto the NCBI Taxonomy (73). The cladogram is rendered using PhyloT v2 and iTOL v6 (75). Animal silhouettes in this and subsequent figures are downloaded from PhyloPic. Other lineages of COL4A found in invertebrates are omitted for clarity and shown in Fig. 4.

**Figure 4.**
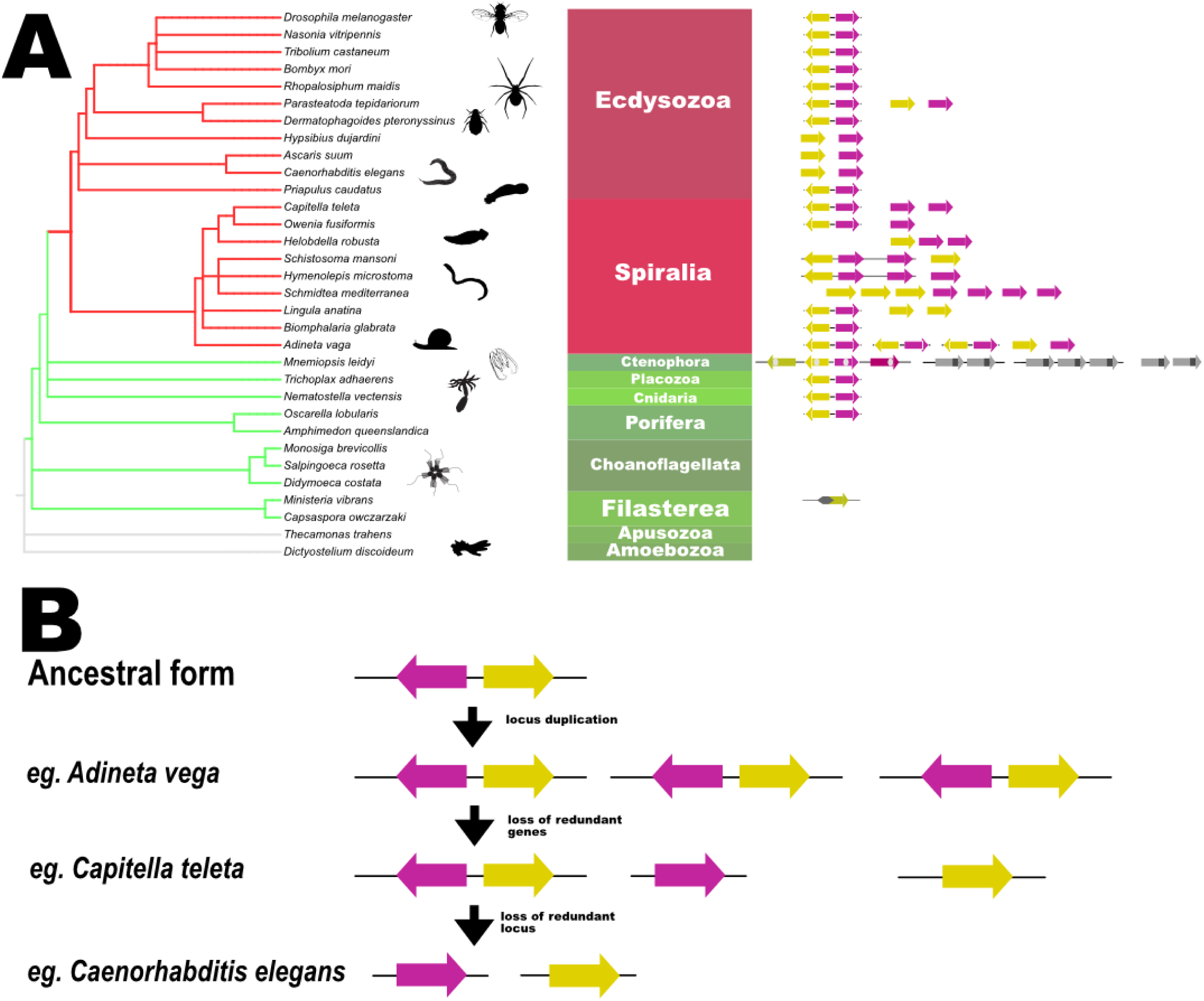
COL4A gene duplications observed in Protostomes and basal metazoans. A. Cladogram of the protostome and basal metazoans showing the diverse gene structure of COL4A genes observed in these animals. Protostomes are in red while the basal metazoans are in green. COL4A1 and COL4A2 derived genes are indicated, where the genes are linked, they are connected with a black line, when unlinked they are separate. Ctenophore COL4A genes have an additional domain and their relationships to COL4A1 and COL4A2 are less clear and indicated in gray. The filasterean *Ministeria vibrans* has a single known COL4A like gene with a distinct N-terminal domain. B. Hypothetical model for the generation of COL4A gene diversity. We propose that COL4A genes may be duplicated as pairs resulting in multiple copies of COL4A gene pairs as in *Adineta vega*. Individual COL4A genes may be deleted for some gene pairs as in *Capitella teleta*. While in other animals the original COL4A⟨1|2⟩ gene pair is lost leaving only the single COL4A1 and COL4A2 genes as is found *C. elegans*.

We propose a plausible path of how diverse COL4A1 and A2 gene arrangements arose in Protostomes (Figure 4B and Supplemental Fig. 1). The gene duplication events, resulting in multiple copies of the COL4 gene pairs as found in *Adineta vega*, may also lead to loss of one of the COL4⟨1|2⟩ pairs either through incomplete gene duplications or accumulation of missense or nonsense alleles in redundant COL4 genes. This may result in the presence of the original COL4A⟨1|2⟩ gene pair and one or more COL4 single copy genes, such as observed in *Capitella teleta*. The COL4A⟨1|2⟩ gene pair could, finally, be lost in some species resulting in two, or more, single copies of the COL4 gene as observed in *C. elegans*. While other evolutionary paths may be possible, this gene duplication hypothesis is consistent with an ancient and conserved COL4A⟨1|2⟩ gene pair, and its ancestral role in the emergence of vertebrate COL4A⟨3|4⟩ and the COL4A⟨5|6⟩ gene-pairs.

We next sought to identify and analyze microsyntenic blocks of the COL4 genes to gain additional evidence for a genetic linkage between COL4A⟨1|2⟩ and COL4A⟨3|4⟩, and a linkage between COL4A⟨1|2⟩ or COL4A⟨3|4⟩ and the COL4A⟨5|6⟩ gene-pair. A microsyntenic block containing each of the three vertebrate COL4 gene-pairs and the IRS gene family was previously noted, wherein an IRS paralog exists in close proximity to each of the three COL4 gene-pair paralogs (Figure 5) (53). This finding prompted us to further identify the order and conservation of genes neighboring the COL4 gene-pairs as an approach for determining genetic linkages.

**Figure 5.**
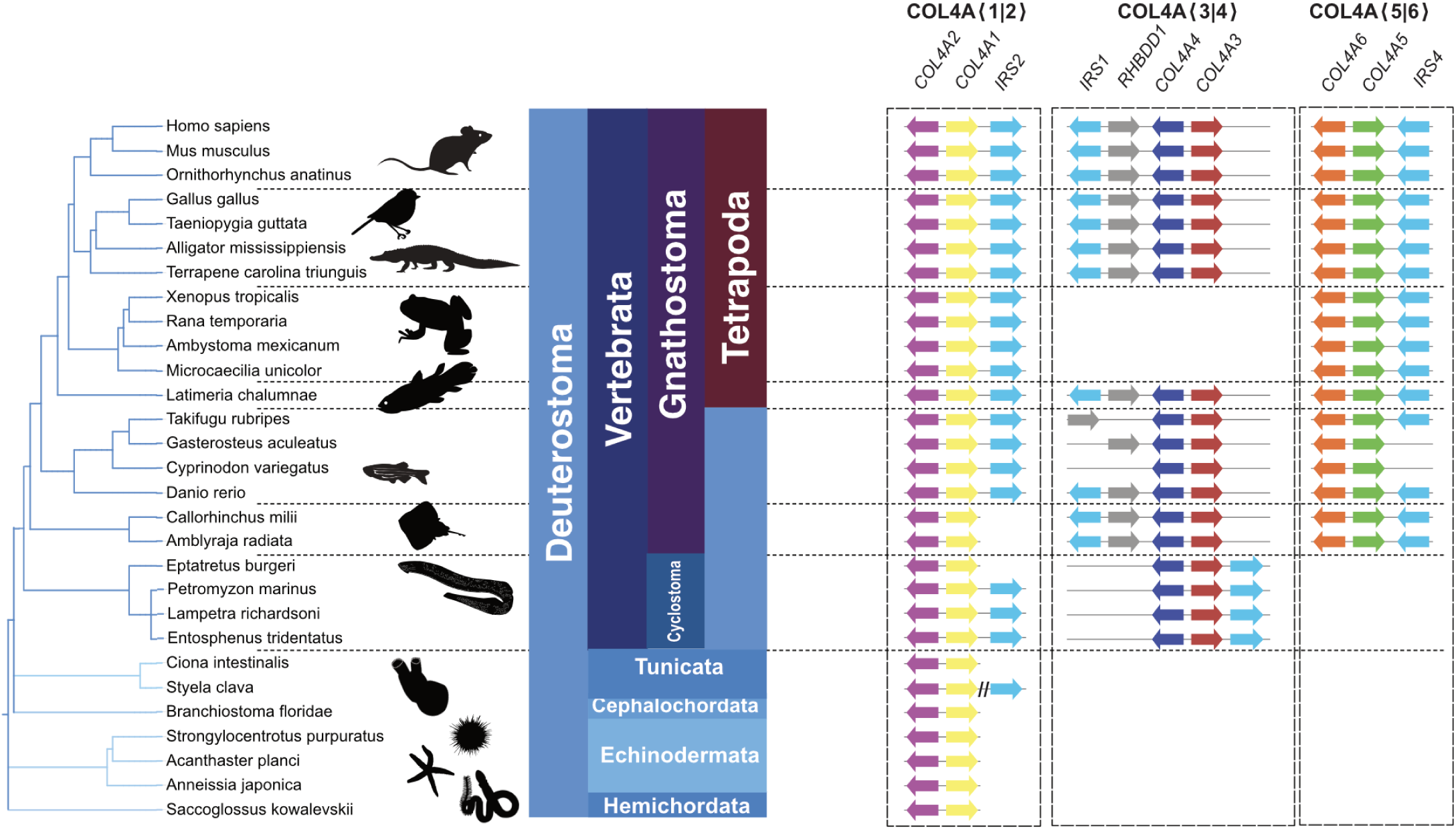
Duplications of the COL4A⟨1|2⟩ gene-pair in Deuterostomes and Vertebrates. Basal vertebrates such as starfish, acorn worms, lancelets and tunicates have a single COL4A⟨1|2⟩ gene pair as found in basal metazoans. In most Deuterostomes the COL4A⟨1|2⟩ gene pair (yellow and magenta) is found tightly linked to an IRS2 ortholog (light blue). Notably, the tunicate *Styela clava* has an IRS ortholog located approximately 14 MBp from the COL4A⟨1|2⟩ gene pair. COL4A⟨3|4⟩ gene pair (red and blue) first appears in cyclostomes and is found in all vertebrate clades except amphibians, often near the IRS1 (light blue) and RHBDD1 (grey) orthologs. The COL4A⟨5|6⟩(orange and green) gene pair is first observed in bony fishes including basal Chondrichthyes and is tightly linked to the IRS4 ortholog (light blue).

### Microsynteny of COL4A⟨1|2⟩ gene-pair in vertebrate evolution

We identified the synteny for the COL4A⟨1|2⟩ gene-pair in diverse species (Figure 6) and found an extensive conservation of neighboring gene order. For example, in all mammals examined, the five genes on either side of the COL4A⟨1|2⟩ gene pair are absolutely conserved in order and orientation. Reptiles have near identical syntenic gene order, while birds have a chromosomal rearrangement 5’ of the MYO16 locus. We next investigated the COL4A⟨1|2⟩ locus in amphibia. Extant amphibia consist of three orders: Salienta (frogs and toads), Caudata (salamanders and newts) and Caecilians. Notably amphibian genomes tend to be very large, making genome assembly and, thus, microsyntenic block analysis difficult due to contig fragmentation. In the salamander genome the COL4A⟨1|2⟩ containing contig is small, limiting microsynteny analysis, however, in the conserved synteny of the COL4A⟨1|2⟩ locus to other vertebrates is clear. The oldest extant group of tetrapods with a well described genome is the Coelacanth whose genome also shows conserved microsynteny, essentially identical to that found in mammals. Thus, in tetrapods the COL4A⟨1|2⟩ microsyntenic block is highly conserved.

**Figure 6.**
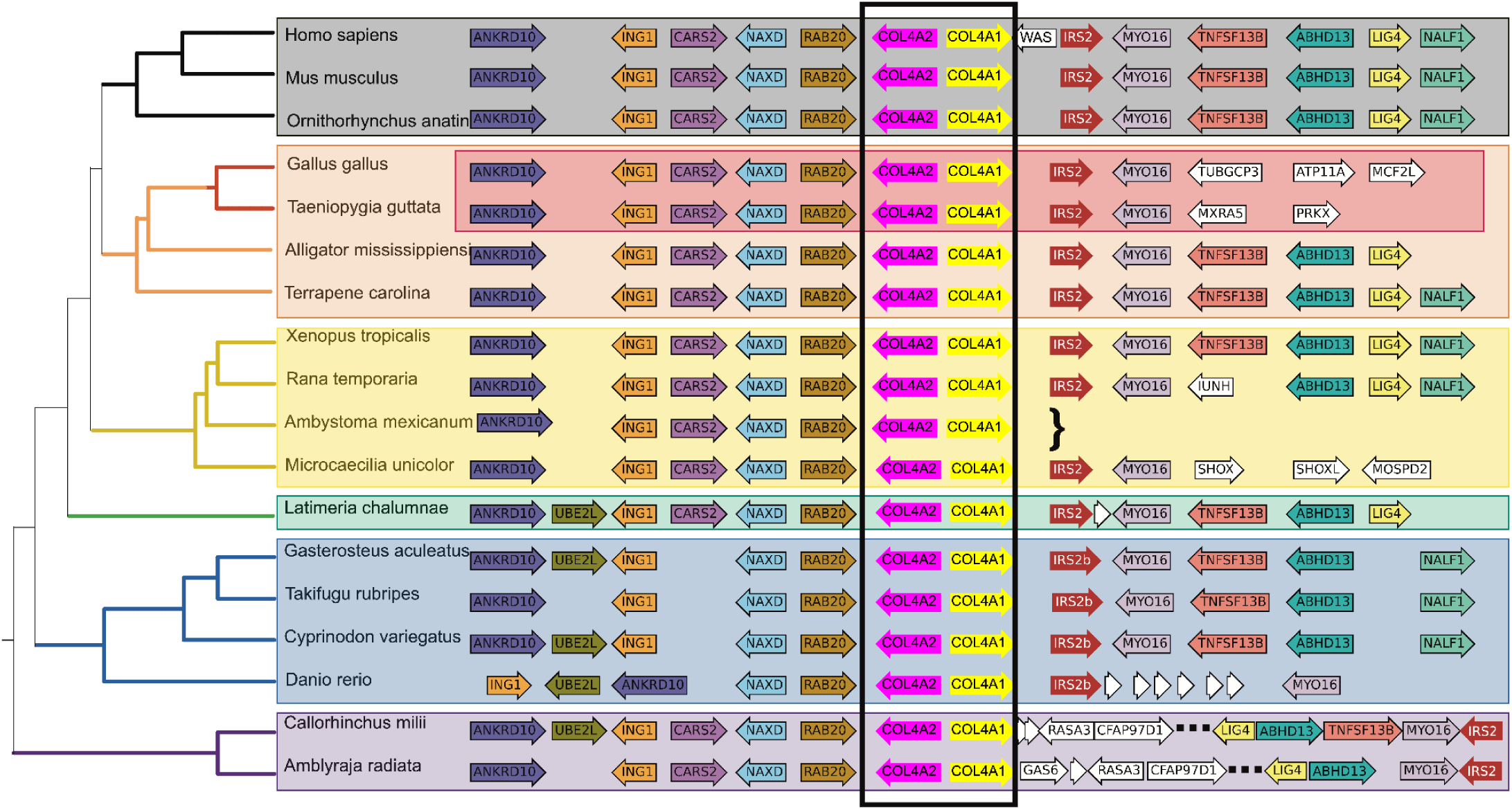
Vertebrate synteny of the COL4A⟨1|2⟩ gene-pair locus. Genome sequences from selected and diverse jawed vertebrates were examined surrounding the COL4A⟨1|2⟩ gene pair and protein coding genes and their orientations were identified. These are indicated as rows next to their gene names for each species. The standard ncbi phylogenetic tree is indicated to the right of each species. The axolotl genome is very large and the contig containing COL4A2 is shown; curly brace (}) marks the boundaries of the contig where the annotated chromosome is likely misassembled. Genes indicated in white are either low confidence gene annotations or genes that are not syntenic to other vertebrates. Note that there is a chromosome inversion in the chondrichthyes fishes. This places the IRS gene distal to the COL4A⟨1|2⟩ locus; the genes immediately adjacent to COL4A⟨1|2⟩ are shown as are the syntenic genes, however intervening genes are omitted for clarity (omitted genes are indicated with the ellipses). The names of genes are based on orthology groups identified through blastp homology to the human genome annotation.

We next investigated whether this synteny was conserved within fishes (Fig. 6). Teleosts, which exhibit extraordinary diversity in ecology, behavior and morphology, may show more diversity which may be more revelatory of the origins of this synteny group, however we found that in diverse teleost groups, the core of the COL4A⟨1|2⟩ microsyntenic region was well conserved. In cartilaginous fishes there is a significant deviation from the microsynteny: the region 3’ of COL4A1 diverged from that observed in other vertebrate groups. However, the microsyntenic block to the region 3’ of COL4A1 occurs nearby on the same chromosome, which suggest that through a chromosomal break and inversion, the microsynteny was lost but synteny was retained. This result suggests that a chromosomal break and inversion proximal to the COL4A1 3’ region occurred distinguishing the Chondrichthyes (cartilaginous fishes) and Osteichthyes (bony fishes and their descendants). Together these findings reveal a highly conserved syntenic block of genes containing the COL4A⟨1|2⟩ gene-pair which was maintained throughout jawed vertebrate evolution.

### Microsyntenies of COL4A⟨3|4⟩ and COL4A⟨5|6⟩ gene-pairs in vertebrate evolution

We identified a microsyntenic region for the COL4A⟨3|4⟩ gene pair and found extensive conservation throughout jawed vertebrate (Gnathostome) evolution (Fig. 7A). Within the tetrapod lineage in mammalian, reptilian and avian genomes, the genes flanking COL4A⟨3|4⟩ are well conserved forming a microsyntenic block (Figure 7A). This conservation is retained in the Coelacanth and cartilaginous fishes, though is less conserved in teleosts. Strikingly, the IRS paralog associated with COL4A⟨3|4⟩ is found in a different orientation and gene order throughout this lineage compared to the COL4A gene pair when compared to COL4A⟨1|2⟩. While IRS2 is found downstream of COL4A1, IRS1 is found downstream of COL4A4 and the RHBDD1 gene intervenes between COL4A4 and IRS1.

**Figure 7.**
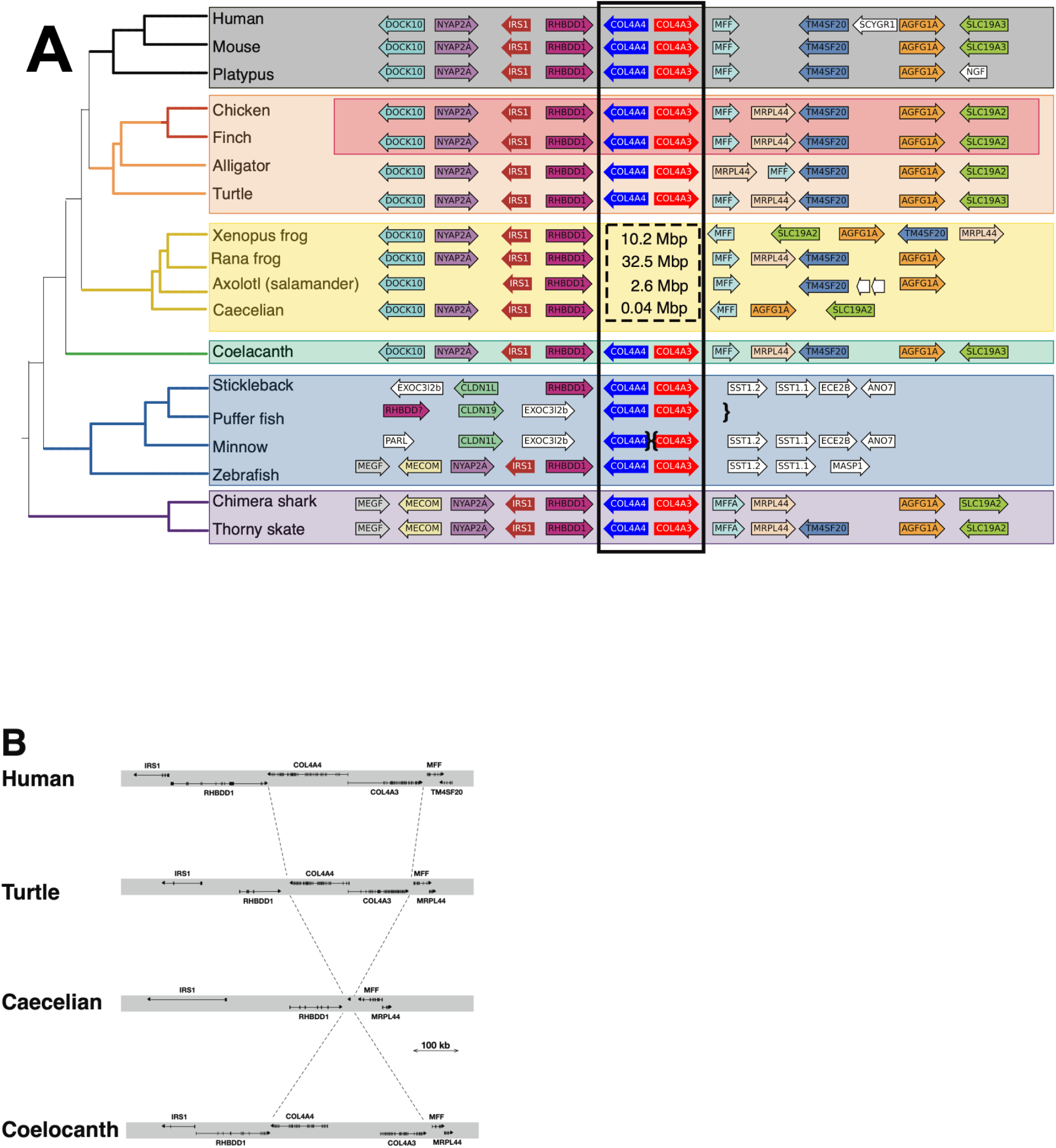
Vertebrate synteny of the COL4A⟨3|4⟩ gene-pair locus, and deletion of the gene-pair in amphibians. A. Genome sequences from selected and diverse jawed vertebrates were examined surrounding the COL4A⟨3|4⟩ gene pair and protein coding genes and their orientations were identified as for Figure 6. In zebrafish, the genes found downstream of COL4A3 in other vertebrates, including SLC19A3, AGFG1, MRPL4, and MFF are found upstream of COL4A1, consistent with a chromosome inversion. The absence of the COL4A⟨3|4⟩ locus in amphibians is indicated with the dashed box. The size of the chromosome fragment in each species replacing the COL4A⟨3|4⟩ locus is indicated in the box. In the Caecilian genome, a single gene encoded by a single exon encoding a kinase similar to TSSK6 in humans is found in this intergenic region, while in Axolotl no genes are observed in the intergenic gap. In the two frog species, multiple genes are found in the intergenic region. B. The exon-intron structure of selected jawed vertebrate genomes reveals the precise deletion of COL4A⟨3|4⟩ in the Caecelian genome: neither neighboring genes are affected save that MFF is inverted in some amphibians.

Surprisingly, the syntenic analyses revealed that the COL4A⟨3|4⟩ gene-pair was deleted in amphibian genomes (Figure 7A). The deletion was precise, retaining the neighboring genes IRS1 and RHBDD1 which are 3’ to COL4A4 in other vertebrates and MFF and MRPL44 which are 3’ to COL4A3 in other vertebrates (Figure 7B). The caecilian genome has a novel single exon protein coding transcript that was detected in the small gap between RHBDD1 and MFF, but there is no clear insertion of functional DNA at this locus compared to those found in both more basal tetrapods (Coelacanth) and more advanced tetrapods (reptiles and mammals). In contrast, larger insertions are observed at the locus in other amphibian classes. Notably, COL4A⟨3|4⟩ gene-pair was absent in other Amphibian orders. Notably, this precise deletion of the COL4A⟨3|4⟩ gene-pair occurs in species that otherwise have quite large and expanded genomes. Parenthetically, this finding of a naturally occurring double gene knockout provided a strategy in a companion paper (25) to gain insights into critical molecular features the Col-IV**^α345^** scaffold that confer function to the GBM.

We next identified the microsyntenic block containing the COL4A⟨5|6⟩ gene-pair. The COL4A⟨5|6⟩ loci in mammalian genomes differs distinctly from both COL4A⟨1|2⟩ and COL4A⟨3|4⟩ which have microsyntenic blocks that are nearly identical (Figure 8). The IRS4 locus is found downstream and encoded by the opposite strand from COL4A5 compared to IRS and COL4A1, suggesting that the duplicated syntenic block which generated these paralogies underwent rapid rearrangements following duplication. The decreased conservation of the syntenic regions surrounding COL4A⟨5|6⟩ is most pronounced in the Teleosts which have quite reduced extents of synteny conservation. Collectively, these results reveal that jawed vertebrates have a well-conserved syntenic block surrounding the COL4A⟨1|2⟩, COL4A⟨3|4⟩ and COL4A⟨5|6⟩ gene-pairs.

**Figure 8.**
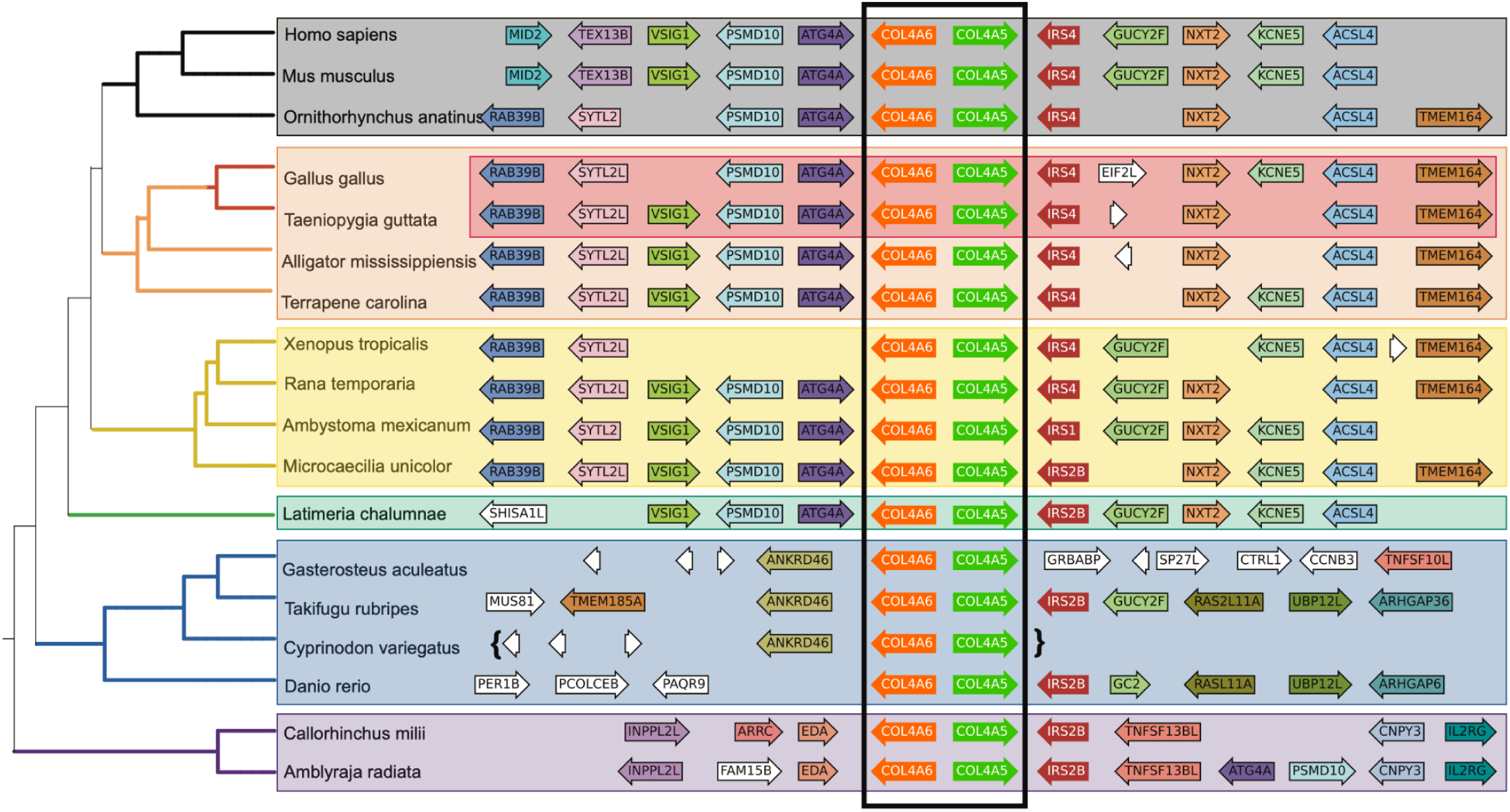
Vertebrate synteny of the COL4A⟨5|6⟩ gene-pair locus. Genome sequences from selected and diverse jawed vertebrates were examined surrounding the COL4A⟨5|6⟩ gene pair and protein coding genes and their orientations were identified as for Figure 2. Note that the synteny is not well maintained in the teleost (blue) lineage perhaps due to the teleostean specific whole genome duplication. Note the presence of the ANKRD46 gene adjacent to COL4A6 in the teleost lineage, which is paralogous to ANKRD10 found downstream of COL4A2, suggesting that these genes share a common ancestry which was duplicated together with the COL4A gene pair generating COL4A⟨5|6⟩.

### Analysis of microsynteny for lineage of COL4 gene-pairs in Deuterostomes

We next sought to analyze the microsyntenic blocks to gain any corroborative evidence for COL4A⟨1|2⟩ to be the progenitor for COL4A⟨3|4⟩ and to determine whether COL4A⟨1|2⟩ or COL4A⟨3|4⟩ is the progenitor of the COL4A⟨5|6⟩ gene-pair. Early Deuterostomes (tunicate, lancelet and starfish) have only a single pair of COL4A genes, corresponding to COL4A⟨1|2⟩ gene-pair and without conserved synteny (Fig. 9). Synteny of COL4A genes first appears in hagfish and lamprey, each of which have two COL 4 gene-pairs, COL4A⟨1|2⟩ and COL4A⟨3|4⟩. In lamprey, both COL4 gene-pairs are adjacent to an IRS paralog, as seen in jawed-vertebrates. However, in hagfish one COL4 gene pair has an IRS paralog adjacent while the other does not. In hagfish, there is an additional synteny conservation in the form of the MYO16 and NYAP2 genes which lie adjacent to each IRS paralog and downstream from the COL4A gene pair. This is a highly similar gene arrangement to those found in the COL4A⟨1|2⟩ and COL4A⟨3|4⟩ synteny groups. This result strongly suggests an evolutionary orthology between the cyclostome COL4 gene pair adjacent to MYO16 and the vertebrate COL4A⟨1|2⟩, as well as a similar relationship between the cyclostome COL4 gene pair which is adjacent to NYAP2 and the vertebrate COL4A⟨3|4⟩. Between the last common ancestor of cyclostomes and jawed vertebrates, the COL4A⟨3|4⟩ gene pair became inverted, moving the NYAP2-IRS genes downstream from COL4A4 rather than downstream from COL4A3 as seen in the cyclostomes. We note also the presence of the genes SLC5A7, FARP2 and BOK downstream of the lamprey COL4A2 gene and STK26, FARP1 and IPO5 downstream of the COL4A4 gene. Paralogs of these genes also appear downstream of COL4A2 and COL4A4 in the vertebrate genome, albeit several Mbp further downstream of the COL4A loci.

**Figure 9.**
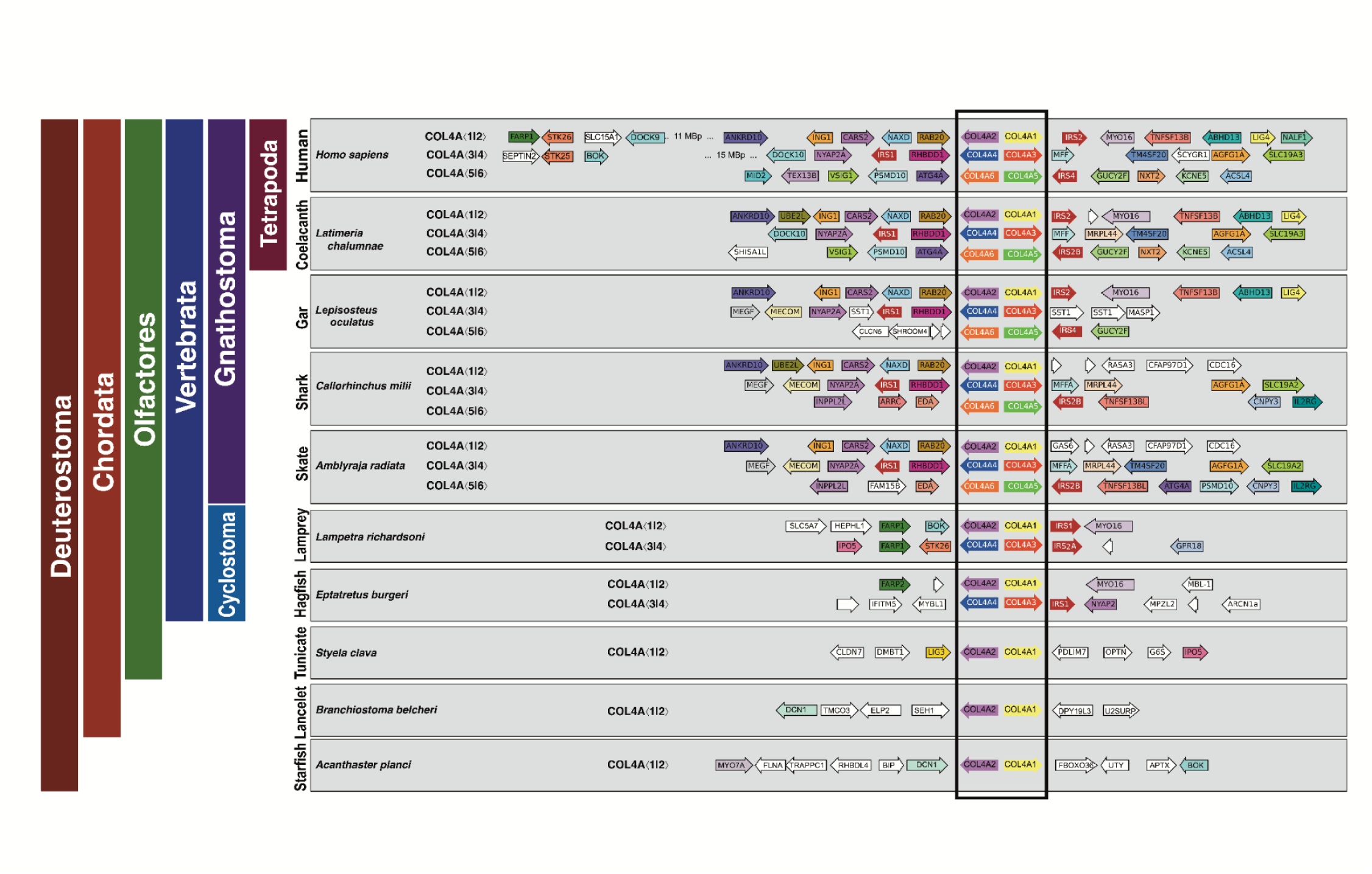
COL4A gene locus evolution in the Deuterostomes. The microsyntenic regions at each COL4A⟨1|2⟩ gene pair (boxed) from selected species are shown. In Deuterostome, the first species to have synteny beyond the COL4A⟨1|2⟩ gene pair is the hagfish which is also the first species to demonstrate duplication of the COL4A⟨1|2⟩ gene pair into COL4A⟨3|4⟩. Lampreys demonstrate increased regional synteny as well as two COL4A gene pairs. Note the presence of paralogs of FARP, BOK, IPO5 and STK25 in lampreys and hagfish that is retained in mammals, albeit at a more distal position on the linkage group. Gnathostomes, except amphibia, show all three COL4A gene pairs.

Collectively, the syntenic analyses reveal that cyclostomes, hagfish and lamprey, have two pairs of COL4 genes, corresponding to COL4A⟨1|2⟩ and COL4A⟨3|4⟩ gene-pairs in vertebrate, and devoid of COL4A⟨5|6⟩. In contrast, shark has six COL4 genes that correspond to vertebrate COL4A⟨1|2⟩, COL4A⟨3|4⟩ and COL4A⟨5|6⟩ gene-pairs. Thus, the COL4A⟨1|2⟩ gene-pair duplicated first in cyclostomes and evolved into the COL4A⟨3|4⟩ gene-pair. Secondarily, in shark a duplication of the COL4A⟨1|2⟩ or the COL4A⟨3|4⟩ gene-pair gave rise to COL4A⟨5|6⟩ in shark; the identity of which pair was not apparent from the microsynteny.

### Analysis of amino acid sequences for lineage of COL4 gene-pairs

We sought to obtain further evidence for lineage of gene duplications from a phylogenetic analysis of the amino acid sequences of the six α-chains, which co-emerged in shark. The analysis for full-length α-chains across metazoans (Fig. 10) and their cognate NC1 domains (Fig. 11), reveal that shark α3 and α5 chains most closely related to the α1 chain. In contrast, the shark α4 and α6 chains related to the α2 chain. These findings suggest that COL4A⟨1|2⟩ is the progenitor for both the COL4A⟨3|4⟩ and COL4A⟨5|6⟩ gene-pairs.

**Figure 10.**
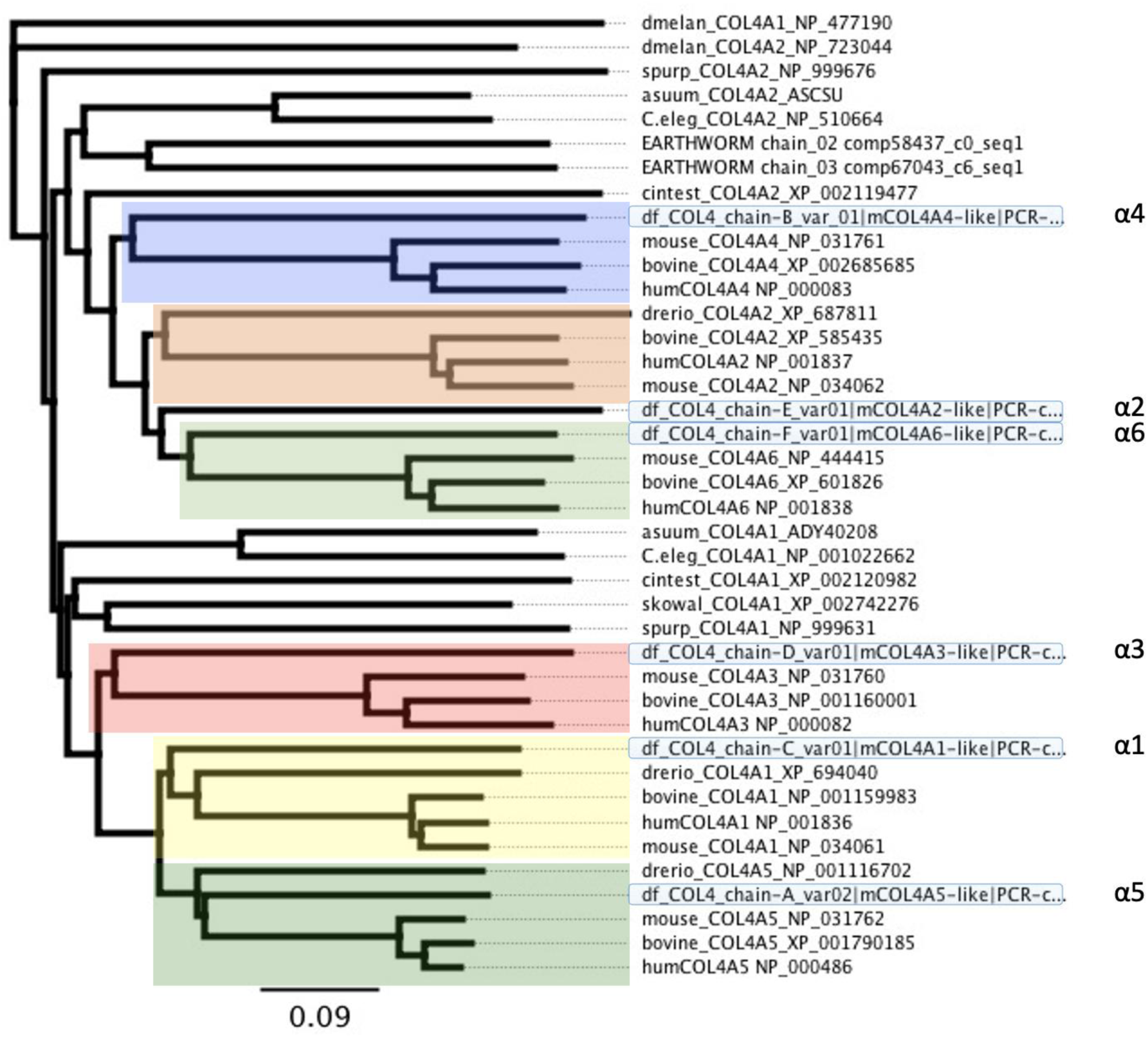
Phylogenetic comparison of Squalus acanthias collagen-IV full-length paralogs to representative metazoan species. Full-length sequences of *S. acanthias* (dogfish) were generated via RNASeq and de novo transcriptome assembly of dogfish basement membrane isolated from kidney, lens. Transcriptome sequences were confirmed by RT-qPCR. Assignment of sequenced dogfish chains was made based on known full length COL4 chains (a1-a6) from human, mouse, bovine, *Drosophila*, zebrafish, *C. elegans*, earthworm, *C. intestinalis*, *S. kowalevsky*, *S. purpuratis*. Phylogenetic tree is built using Neighbor-Joining method using Geneious Bioinformatic Software (https://www.geneious.com/).

**Figure 11.**
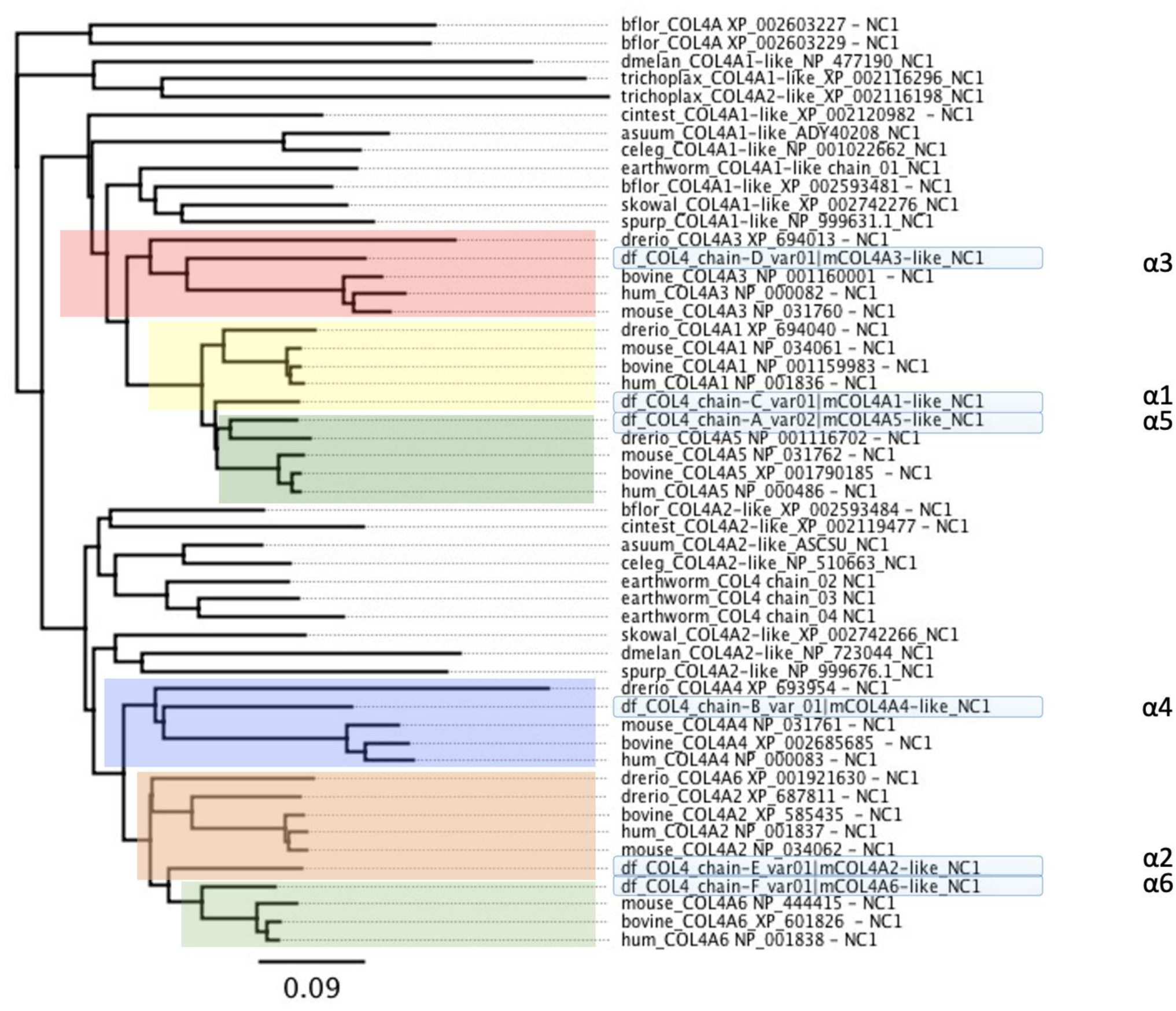
Phylogenetic comparison of Squalus acanthias collagen-IV NC1-domain paralogs to representative metazoan species. Full-length sequences of S. acanthias (dogfish) were generated via RNASeq and de novo transcriptome assembly of dogfish basement membrane isolated from kidney, lens. Transcriptome sequences were confirmed by RT-qPCR. Assignment of dogfish NC1-domains from RNASeq derived full-length chains was made based on known COL4 NC1-domains (a1-a6) from human, mouse, bovine, Drosophila, zebrafish, C. elegans, earthworm, C. intestinalis, S. kowalevsky, S. purpuratis. Phylogenetic tree is built using Neighbor-Joining method using Geneious Bioinformatic Software (https://www.geneious.com/).

### Expression of COL4 gene-pairs in hagfish and shark kidneys

The conservation of COL4 gene-pairs across metazoans posits the question of whether the COL4A⟨1|2⟩ and COL4A⟨3|4⟩ gene-pairs in hagfish and the COL4A⟨1|2⟩, COL4A⟨3|4⟩ and COL4A⟨5|6⟩ gene-pairs in shark are expressed and incorporated into the kidney. At the time this study began, genomic data for these species were unavailable. Therefore, to probe this question, we isolated total RNA from both hagfish and dogfish (shark) kidneys and performed next generation RNA sequencing. De novo assembly of the dogfish transcriptome was performed and used to design specific primers to PCR COL4A transcripts that were subsequently sequenced to confirm the transcriptomics results. We found that at least three COL4A transcripts (corresponding at the protein level to the Col-IV α1, α2, and α3 or α4 chains) were expressed in hagfish kidney, whereas all six transcripts (corresponding to the α1 to α6 chains) were expressed in dogfish kidney (Supplemental text).

We determined whether the Col-IV α1-α6 chains were incorporated into hagfish and dogfish kidney in the form of supramolecular scaffolds. We used the well-established method of characterization of the non-collagenous NC1 hexamers from the scaffolds, as direct evidence for scaffold organization (54). NC1 hexamers were excised by collagenase digestion, purified by size exclusion chromatography, and characterized by SDS/PAGE. The electrophoresis patterns were similar to that of mammalian hexamer by the presence of NC1 dimer and monomer subunits, revealing scaffold expression in both hagfish and shark (29,55) (Fig. 12A).

**Figure 12.**
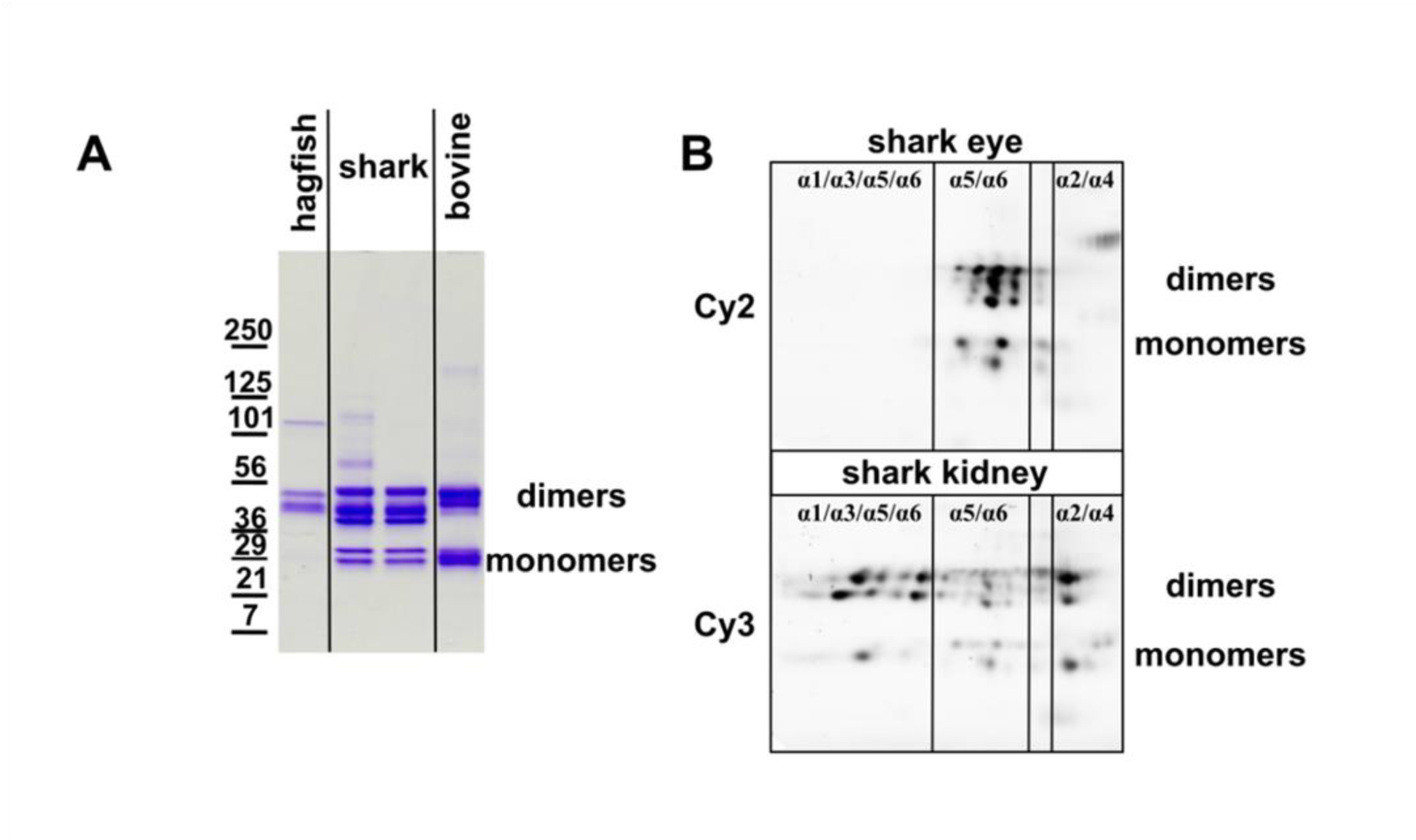
Tissue specific collagen IV chain composition. A. SDS Polyacrylamide Gel Electrophoresis of purified NC1 domains of collagen IV derived from hagfish kidney, shark ocular lens and kidney, and bovine kidney. The shark and bovine NC1 domains run as both uncrosslinked monomers and crosslinked dimers while the hagfish protein does not have detectable monomers. The shark dimer band runs as a set of three bands likely reflecting scaffolds of different compositions. B. Fluorescent images of the 2D-NEPHGE gel loaded with purified NC1 domains from the shark ocular lens (Cy2) and kidney (Cy3). Interestingly, shark ocular lens consists primarily of α5 and α6 while shark kidney has all chains present.

To determine which α-chains were incorporated into shark kidney, we identified the α-chain origin of the NC1 hexamer subunits. The analysis was performed by using two-dimensional non-equilibrium pH gel electrophoresis (2D-NEPHGE) on kidney and lens tissues. Kidney hexamer was labeled with the Cy3 fluorescent dye and lens hexamer with Cy2. The fluorescent-labeled hexamers were mixed and resolved by 2D-NEPHGE (Figure 12B). The lens and kidney patterns were distinct, revealing tissue specific differences in the expression of α-chains. Individual spots on the gels were excised and analyzed with high resolution liquid chromatography and tandem mass spectrometry (LC-MS/MS). A custom dogfish protein database, based on the transcriptomics data, was used to identify α-chain identity of each spot, Supplemental Figure 1, Figure 12). The results show that all six Col-IV α-chains (α1 through α6) were expressed in shark kidney, whereas only the α5 and α6 chains in lens. The scaffold compositions of these multiple α-chains remain unknown, but they do not include the Col-IV**^α345^** scaffold in shark kidney as described in a companion paper (25).

### Evolutionary changes in the number of cysteine codons in the COL4A⟨3|4⟩ gene-pair

In prior studies, we found that Col-IV**^α345^,** the principal component of GBM, is highly crosslinked by disulfide bonds in comparison to the Col-IV**^α121^** scaffold (56). Specifically, the number of cysteine residues is very large in the α3 and α4 chains, in comparison to the other Col-IV α1, α2, α5 and α6-chains. We speculated that disulfide crosslinks are a key structural feature that confers mechanical strength to the GBM, as well as to protect against proteolysis (56). To gain insights into the role of cysteine residues, we investigated the phylogenetic distribution of their codons in the COL4 gene-pairs.

We identified a significant increase in the cysteine codons in the COL4A⟨3|4⟩ gene pair, compared to COL4A⟨1|2⟩ and COL4A⟨5|6⟩ (Figure 13A), and also a very large variation in numbers of codons among diverse species (Figure 13B). The variance occurred both within and between clades (Figure 14A), and specifically when vertebrates were classified by their environment rather than their clade (Figure 14B). A statistically significant difference was identified within the teleost group when they were classified by the salinity of their environment (Figure 14B). Fish in freshwater environments have increased cysteine content compared to those found in estuarine and marine environments. Notably however, this difference in cysteine content is not present in the cyclostomes (Fig. 13A).

**Figure 13.**
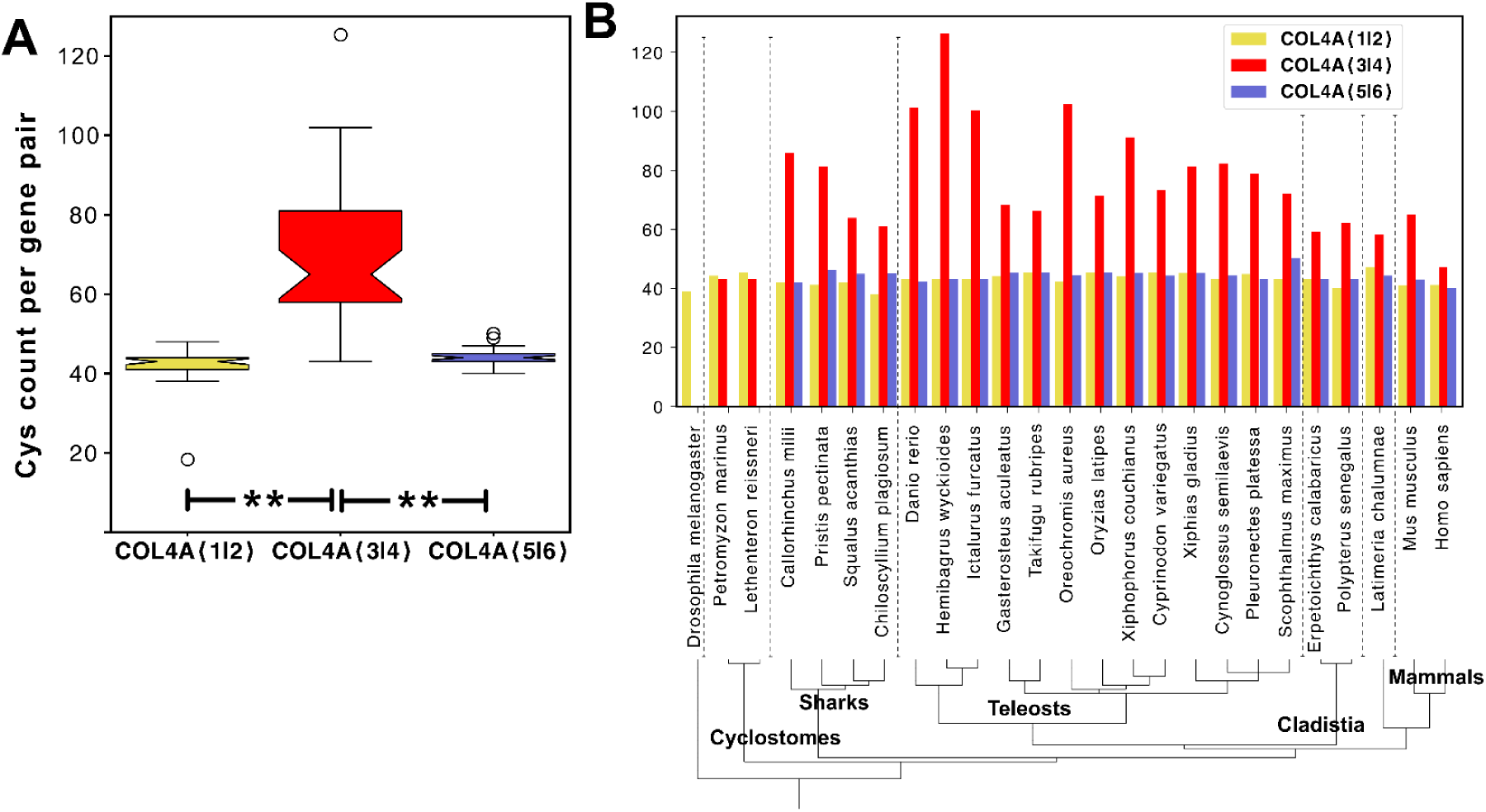
Cysteine content is increased in COL4A⟨3|4⟩ gene-pairs. A. Both COL4A⟨1|2⟩ and COL4A⟨5|6⟩ encode similar numbers of cysteine residues per gene pair while the COL4A⟨3|4⟩ gene pair encodes an increased number of cysteine residues in diverse vertebrates. B. Comparison of the cysteine codon count per gene pair in diverse vertebrate species is not correlated with clade.

**Figure 14.**
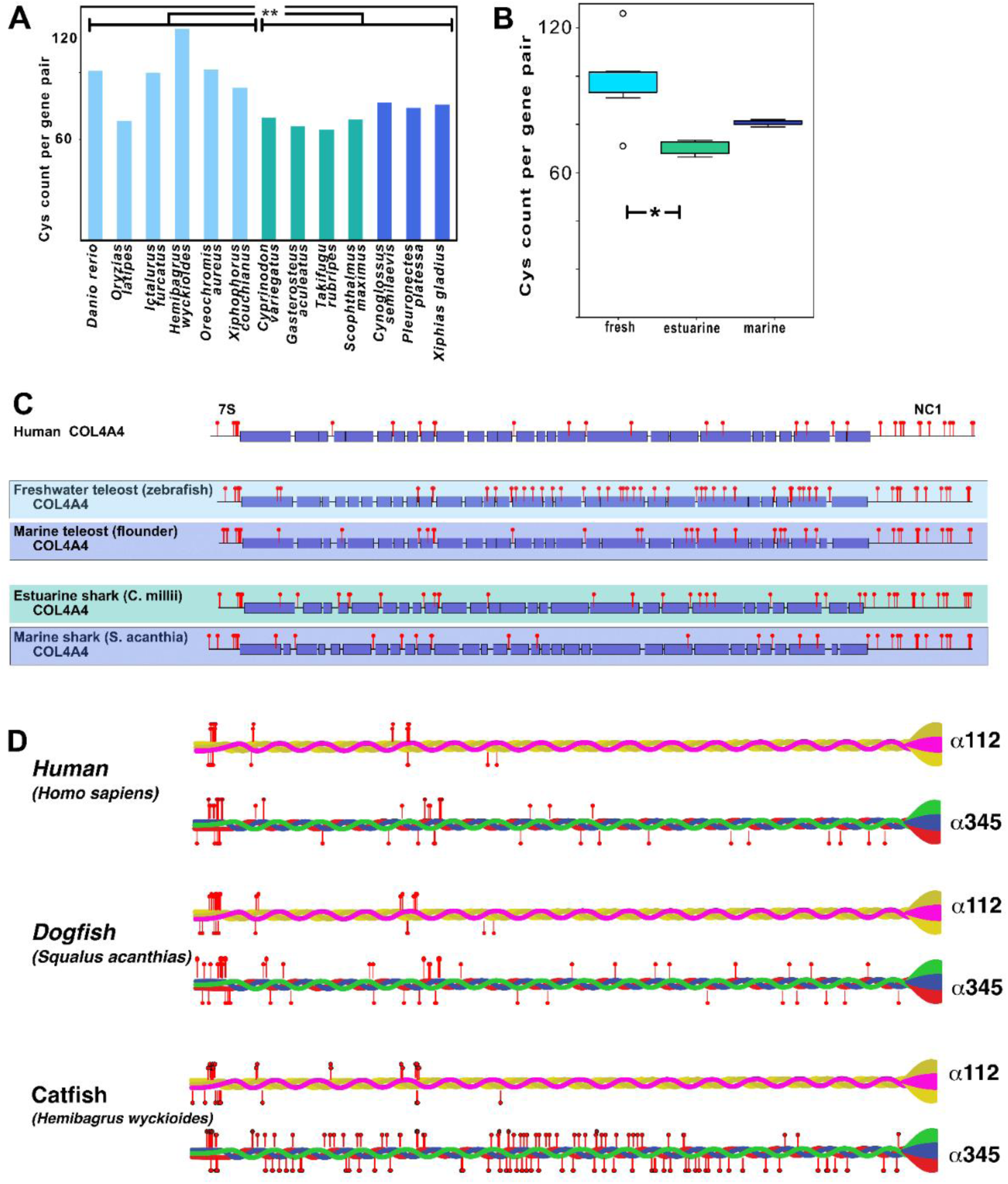
Cysteine content in COL4A⟨3|4⟩ gene-pair varies by animal habitat. A. Cysteine count in teleosts found in different osmolarity habitats (freshwater, pale blue; estuarine, teal; and marine, dark blue). Freshwater fishes have increased cysteine content compared to marine and estuarine fishes. B. Cysteine count in diverse fishes including teleosts and cartilaginous fishes confirming the increased cysteine counts in freshwater fishes generally. C. Map of the cysteines (red pins) on the COL4A4 reading frame from humans and diverse fishes. Collagen encoding domains are colored boxes and non-collagenous domains are indicated by lines, 7S and NC1 domains are indicated at the N termini and C termini respectively. D. Cysteines mapped to the protomer for human, estuarine shark and freshwater catfish Col-IV**^α345^**. Odd numbered COL4A genes are mapped above and even numbered COL4A genes below the protomer schematic. Cysteines in the NC1 domain are known to form disulfide bonds that stabilize the domain and are not shown (78,79).

We next determined whether there was a pattern to the change in cysteine abundance that may relate to function. When the cysteine codons were plotted on the predicted open reading frame of the COL4A4 gene, we found that the variable cysteine residues were scattered throughout the region of the collagenous domain (Figure 14C, and Supplemental Figure 2). The increase in cysteine content was most notable in the C-terminal half of the collagenous domain. At the protein level, the abundance of cysteine residues is a distinguishing feature of the triple-helical protomers that assemble into the Col-IV**^α345^** scaffold, in comparison with the Col-IV**^α121^** scaffold (Fig. 14D). Notably, catfish has the largest number of cysteine residues in a Col-IV**^α345^** protomer, suggesting a key adaptation enabling GBM function in freshwater animals.

## Discussion

In our previous work, we found that collagen IV is a primordial component of basement membranes that enabled the assembly of a fundamental architectural unit for the genesis and evolution of multicellular tissues (1). Also, we found that the structural domains of vertebrate collagen IV protomers, described in Fig. 2, are conserved across metazoans. Moreover, the pairwise arrangement of COL4 genes also appeared to be conserved, based on the analysis of a limited number of species, but the phylogeny and identity of gene-pairs remain unknown.

Here, we sought to extend these phylogenetic analyses to determine the emergence and genetic lineage of the COL4 gene family, with an emphasis on those encoding the Col-IV**^α345^** scaffold. We anticipated this strategy would provide insights into structure-function relationships of the Col-IV**^α345^** scaffold that enabled the GBM to function as an ultrafilter of proteins. We found that the COL4A⟨1|2⟩ gene-pair emerged in basal Ctenophores and Cnidaria phyla and is highly conserved across metazoans (Fig. 15). The COL4A⟨1|2⟩ duplicated and arose as the progenitor to the COL4A⟨3|4⟩ gene-pair in cyclostomes, coinciding with emergence of kidney GBM, and expressed and conserved in jawed-vertebrates, except for amphibians, and a second duplication as the progenitor to the COL4A⟨5|6⟩ gene-pair and conserved in jawed-vertebrates. These findings revealed the genetic emergence and expression of the Col-IV α1 and α2 chains that assemble into the Col-IV**^α121^** scaffold and incorporate ubiquitously in metazoan basement membranes. Moreover, Col-IV**^α121^** is the progenitor scaffold that evolved into vertebrate Col-IV**^α345^** scaffold and incorporated mainly in the GBM of mammals (Fig. 16). In a companion paper, we found that the emergence of the Col-IV**^α345^** scaffold enabled the assembly of a compact GBM that functions as the primary ultrafilter of proteins in mammals (25) (Elena).

**Figure 15.**
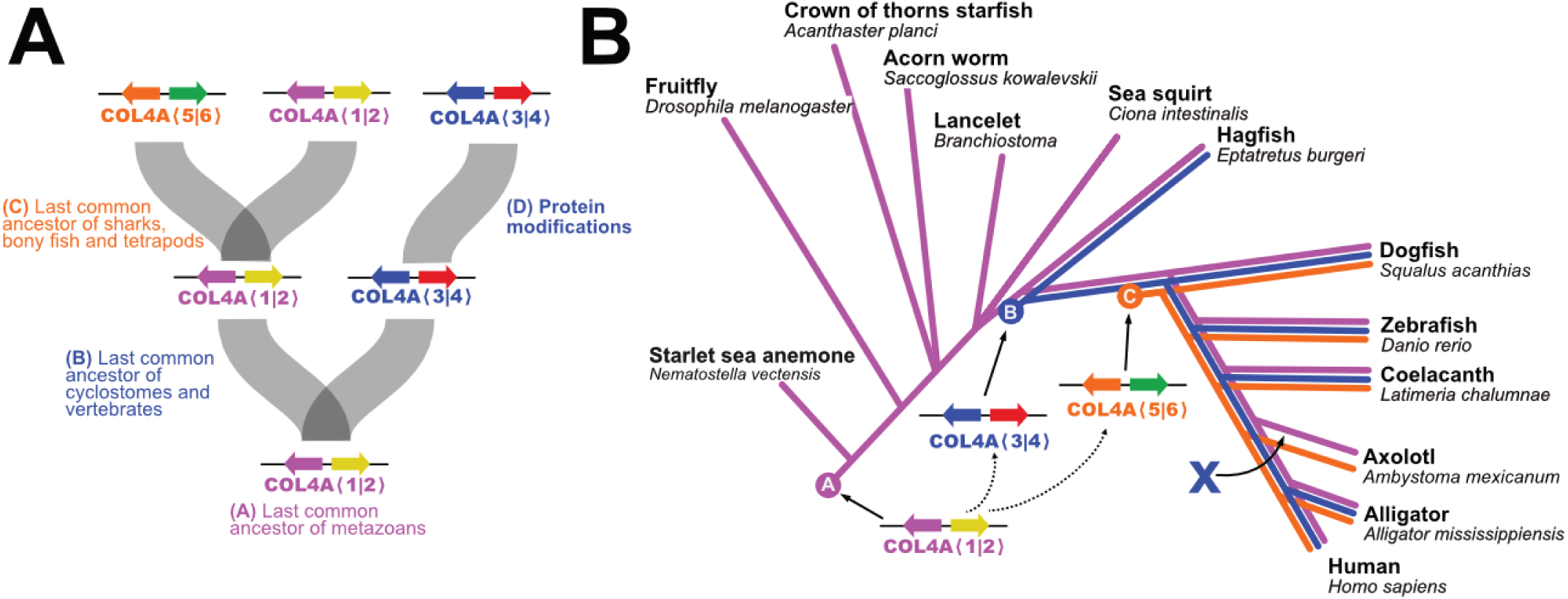
COL4A gene evolutionary history in multicellular animals. A. Hypothesized gene duplication events resulting in generation of the COL4A⟨3|4⟩ and COL4A⟨5|6⟩ with the apparent last common ancestors at each step indicated. B. Highly schematic cladogram of each duplication event (indicated by the letters A, B and C) and each gene pair indicated by the color of the gene pair. The X indicates the loss of the COL4A⟨3|4⟩ gene pair observed in extant Amphibians.

**Figure 16.**
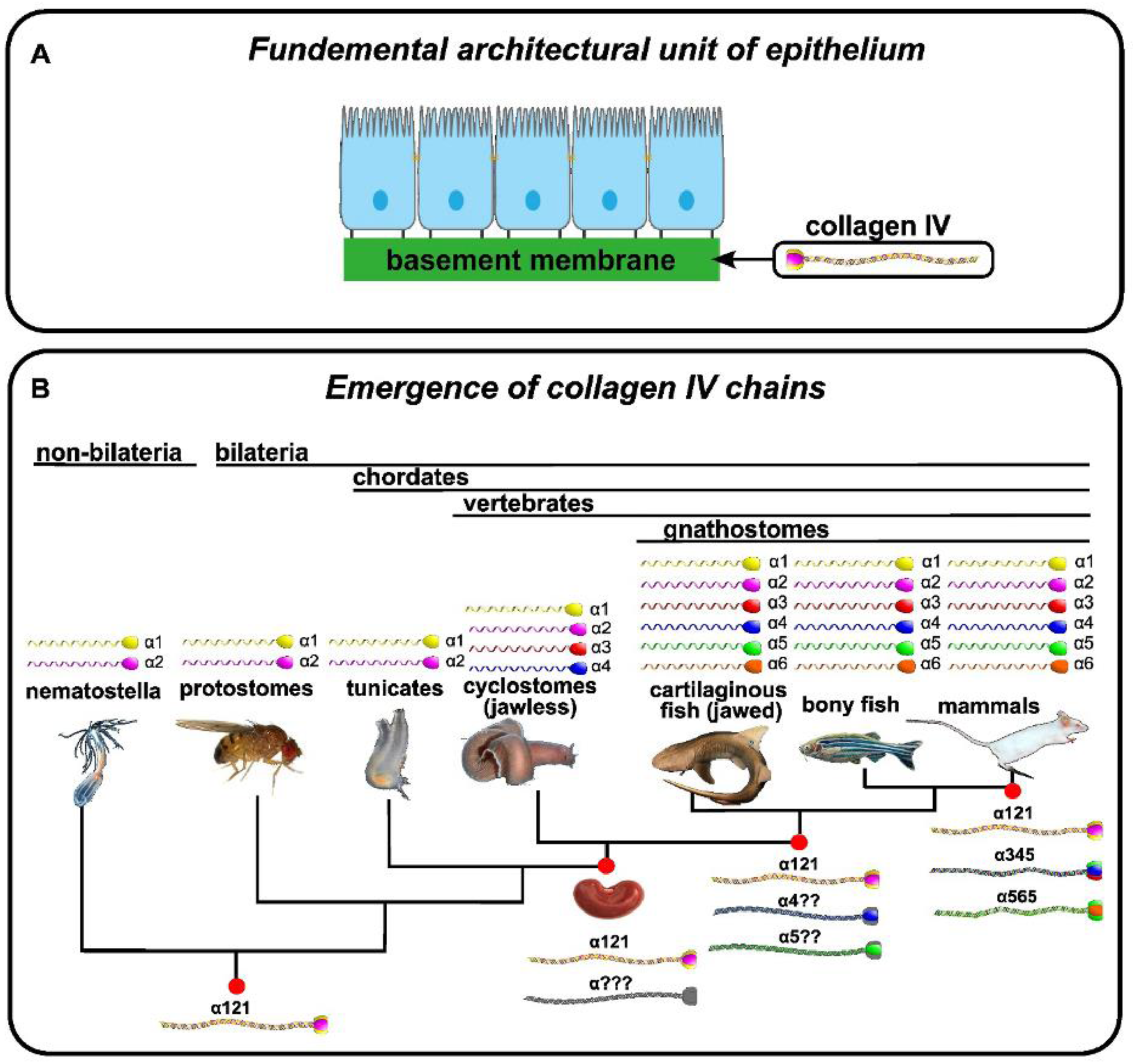
Evolutionary emergence of collagen IV α3 through α6 chains coincides with kidney appearance followed by GBM morphological transition upon emergence of Col-IV^α345^. A. This architectural unit of epithelia tissues is characterized by a layer of apical/basal-polarized cells that are laterally connected by tight junctions between plasma membranes, which are basally anchored via integrin receptors embedded in plasma membranes to a basement membrane supra-scaffold. In turn, this architectural unit served as the building block that enabled the formation and evolution of epithelial tissues, the ever-increasing complexity and size of organisms, and for the expansion and diversity of the animal kingdom. Col-IV triple helical protomers, a principal component of basement membranes, was a primordial innovation in early metazoan evolution that enabled the transition to multicellularity and the evolution of epithelial tissues in metazoa (1). B. Genome duplication events led to appearance of COL4A⟨3|4⟩ and then COL4A⟨5|6⟩. Animals have two or more chains of collagen IV generating the Col-IV**^α121^** scaffold. In cyclostomes the α3 and α4 chains first appeared; and α5 and α6 appeared later in cartilaginous and bony fishes. Emergence of COL4A⟨3|4⟩ coincides with the appearance of the glomerulus which generates a high volume of filtrate which is processed by the nephron tubule. However, the mammalian GBM morphology and ultrafilter function are only found in vertebrates that evolved to have Col-IV^α345^ scaffold (25).

In prior studies, we found that Col-IV**^α345^,** the principal component of GBM, is highly crosslinked by disulfide bonds in comparison to the Col-IV**^α121^** scaffold (Fig. 17) (56). We speculated that disulfide crosslinks are a key structural feature that confers mechanical strength to the GBM, as well as to protect again proteolysis. Intriguingly, in the present study, we found that the Col-IV**^α345^** scaffold has an increased number of cysteine residues in comparison to Col-IV**^α121^**, which mediate disulfide crosslinks between protomers, and which vary in number with the osmolarity of the environment. The GBM is the only known basement membrane in which there is bulk flow of liquid rather than diffusion across the membrane. This bulk flow, associated with glomerular ultrafiltration, places high hydrostatic pressure (57) on the GBM. In freshwater teleost, such as catfish and zebrafish living in low osmolarity environments, there is high liquid flux across the GBM, compared to marine animals in high saline environments. Thus, the increase in cysteine content likely confers additional mechanical strength to GBM to withstand high hydrostatic pressure. This environmental adaptation in disulfide crosslinking pinpoints a role for Col-IV**^α345^** scaffold in the GBM versus Col-IV**^α121^**, wherein Col-IV**^α345^** having an increased number of crosslinks enabled the assembly of a compact GBM that withstands the high hydrostatic pressure in mammals (Fig. 17) (25).

**Figure 17.**
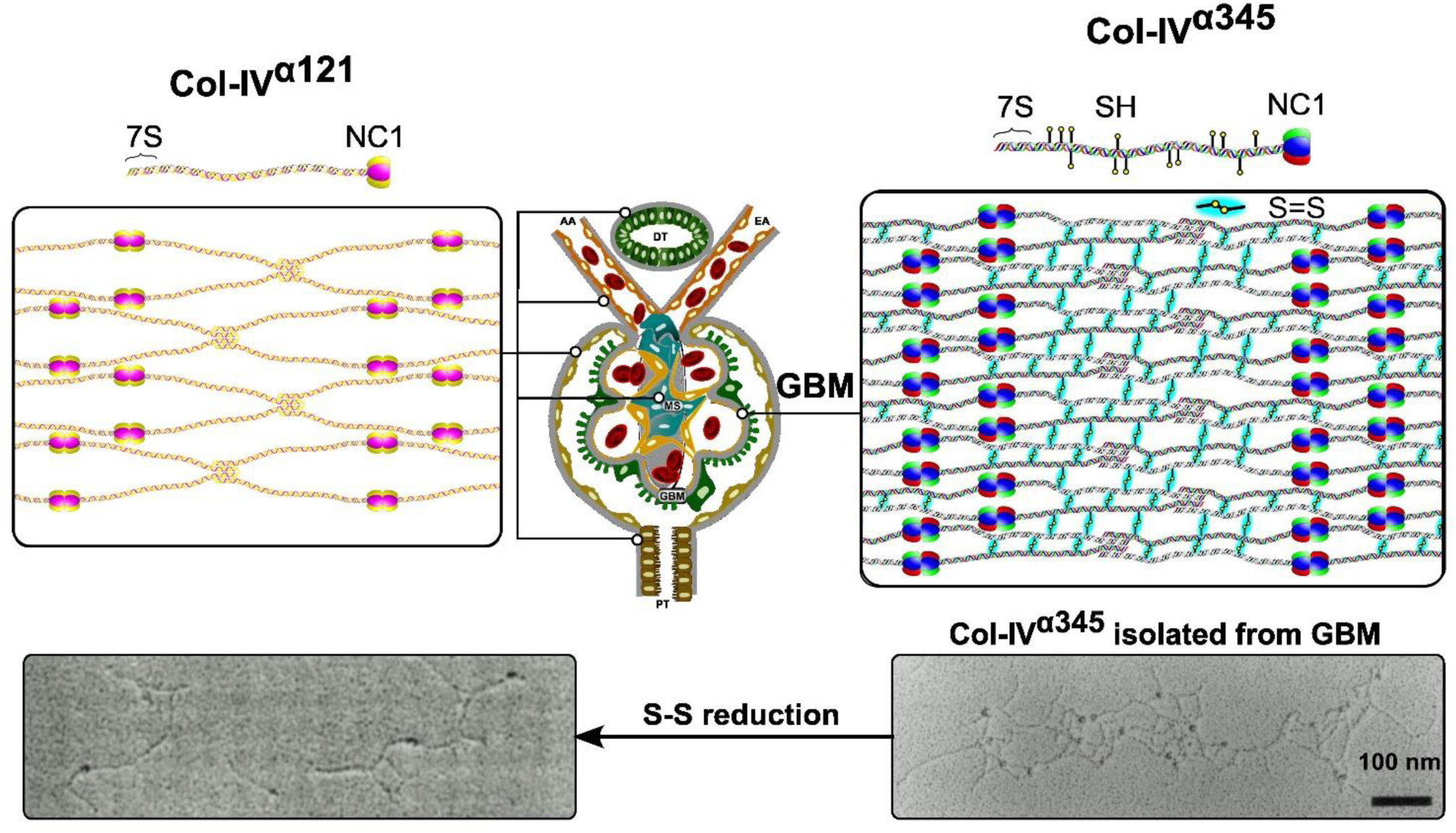
Reinforcement of collagen IV^α345^ scaffolds in the mammalian GBM by disulfide-mediated crosslinks between protomers. A. The Col-IV**^α121^** scaffold schematically represented on the left is a component of the basement membranes surrounding kidney tubules (PT), efferent and afferent arterioles (EA and AA) and Bowman’s capsule. It is also found in the mesangial space (MS). The Col-IV**^α345^** scaffold shown on the right is the principal component of the GBM. This scaffold is reinforced by multiple disulfide bonds (S-S bonds highlighted in cyan) between protomers. B. Rotary shadowing electron microscopy images of Col-IV**^α345^** scaffold, isolated from bovine kidney glomeruli (right), and the same scaffold after reduction of the disulfide bonds (left). Upon reduction of lateral disulfide crosslinks, the scaffold dissociates into protomers which are dimerized through their NC1 domains (56).

It is noteworthy that the GBM of terrestrial amphibians is devoid of the Col-IV**^α345^** scaffold (25). The terrestrial environment, rather than freshwater, required quite distinct water retention mechanisms, suggesting that Col-IV**^α345^** scaffold was counterproductive. Unlike most vertebrates in which the glomerular filtrate drains from Bowman’s space into the proximal tubule, the amphibian glomerular filtrate drains into the coelom (58) and the coelomic fluid then passes into the nephron tubule. Such an anatomical arrangement increases the surface area available for protein and electrolyte absorption, perhaps obviating the need for high fluid flow across the GBM and the necessity for a compact GBM. Instead, amphibians adapted the slit diaphragm as the primary ultrafilter of proteins.

### Experimental Procedures

#### Next Generation RNA Sequencing

Transcriptomes used in this study were sequenced at the Vanderbilt Technologies for Advanced Genomics Core Facility (VANTAGE, Nashville, TN). The Illumina TruSeq mRNA Sample Preparation Kit was used to convert the mRNA in 100 ng of total RNA into a library of template molecules suitable for subsequent cluster generation and sequencing on the Illumina HiSeq 2500 using the rapid run setting. The pipeline established in VANTAGE was followed and is briefly described below. The first step was a quality check of the input total RNA by running an aliquot on the Agilent Bioanalyzer to confirm RNA integrity. The Qubit RNA fluorometry assay was used to measure sample concentrations. The input-to-library prep was 100 ng of total RNA (2 ng/ul). The poly-A containing mRNA molecules were concentrated using poly-T oligo-attached magnetic beads. Following purification, the eluted poly(A) RNA was cleaved into small fragments of 120–210 base pair (bp) using divalent cations under elevated temperature. The cleaved RNA fragments were copied into first strand cDNA using SuperScript II reverse transcriptase and random primers. This step was followed by second strand cDNA synthesis using DNA Polymerase I and RNase H treatment. The cDNA fragments then went through an end repair process, the addition of a single ‘A’ base, and then ligation of the Illumina multiplexing adapters. The products were then purified and enriched with PCR to create the final cDNA sequencing library. The cDNA library then undergoes quality control by running on the Agilent Bioanalyzer HS DNA assay to confirm the final library size and on the Agilent Mx3005P qPCR machine using the KAPA Illumina library quantification kit to determine concentration. A 2 nM stock was created and samples pooled by molarity for multiplexing. From the pool, 12 pmoles were loaded into each well for the flow cell on the Illumina cBot for cluster generation. The flow cell was then loaded onto the Illumina HiSeq 2500 utilizing v3 chemistry and HTA 1.8. The raw sequencing reads were processed through CASAVA-1.8.2 for FASTQ conversion and demultiplexing. The Illumina chastity filter was used and only the PF (passfilter) reads are retained for further analysis. *De novo* assembly of transcriptomes was performed using Velvet/Oases, and Trinity software packages with default settings (59–61). The accuracy of *de novo* assembly was checked in a parallel next generation RNA-Seq experiment using RNA from mouse PFHR9 cells. De novo assembled transcripts were used to generate BLAST databases to search for Collagen IV hits using tblastn (62) with e-value cutoff set to 10^−15^. Multiple sequence alignments and conserved domain searches were performed with the Geneious v5-6 software (Biomatters).

#### Cloning of Hagfish and Dogfish COL4A

To confirm accuracy of hagfish and dogfish COL4A cDNA sequences obtained from RNAseq-based sequences, we performed a series of RT-PCR cloning experiments using primers designed to NGS-detected COL4A candidates. RNA was prepared using the QIAGEN One-Step RT-PCR Kit and PCR products were cloned using the QIAGEN PCR Cloning Kit.

#### Isolation, purification, and analysis of Collagen IV NC1 hexamers

Tissues were frozen in liquid nitrogen, pulverized in a mortar and pestle and then homogenized in 2. 0 ml g^−1^ digestion buffer and 0. 1 mg ml^−1^ Worthington Biochemical bacterial collagenase and allowed to digest at 37°C, with spinning for 24 hr. Liquid chromatography purification of solubilized NC1 varied by species based on protein yield. All ctenophore NC1s were purified by gel-exclusion chromatography (GE Superdex 200 10/300 GL). For reduction and alkylation of Collagen IV NC1 hexamers, fractions containing high-molecular-weight complex from size-exclusion chromatography were concentrated by ultrafiltration and reduced in TBS buffer with various concentrations of DTT. After incubation for 30 min at 37°C, samples were alkylated with twofold molar excess of iodoacetamide for 30 min at room temperature in the dark. After mixing with SDS loading buffer, samples were heated for 5 min in a boiling water bath and analyzed by non-reducing SDS-PAGE. Collagenase-solubilized NC1 hexamers were analyzed by one-dimensional SDS-PAGE in 12% *bis*-acrylamide mini-cells with Tris-Glycine-SDS running buffer. Sample buffer was 62. 5 mM Tris-HCl, pH 6. 8, 2% SDS (w/v), 25% glycerol (w/v), 0. 01% bromophenol blue (w/v). Western blotting of SDS-dissociated NC1 hexamer was developed with JK-2, rat monoclonal antibody (kindly provided by Dr. Yoshikazu Sado, Shigei Medical Research Institute, Okayama, Japan). All Western blotting was done with Thermo-Scientific SuperSignal West Femto chemiluminescent substrate and digitally imaged with a Bio-Rad GelDoc system. Two-dimensional NEPHGE electrophoresis was performed according to the original protocol developed (63) with slight modifications developed in Hudson lab and used successfully to separate NC1 domains of Collagen IV (64).

#### Proteomics

All proteomics experiments were done at Vanderbilt’s MSRC Proteomics Core facility. Major protein spots from Coomassie stained 2D-NEPHGE separated dogfish Collagen IV preparations isolated from various tissues were cut out of the gel and proteins were identified and LFQ-quantified using MaxQuant (65,66). The heatmap was created with heatmapper. ca based on LFQ-data (67).

#### Synteny analysis

Well-characterized genomes were selected from those appearing in the NCBI and Ensembl databases (68,69) and the Axolotl genome (https://genome.axolotl-omics.org/cgi-bin/hgGateway; assembly ambMex 6. 0-DD) and COL4 gene sequences were identified based on text and blastp (70) searches. Syntenic genes were then manually identified and their orientation and order on the chromosome annotated and plotted with matplotlib (71). Protein domains were mapped to their ORFs using the Conserved Domain Database (72). The COL4 gene phylogeny was derived from the NCBI Taxonomy (73,74) and plotted using iTOL (75).

## Acknowledgments

We wish to thank Calvin Greenwood Beames Jr. who told BGH that the intestinal basement membrane of the nematode *Ascaris* was visible in low power microscopes in 1973 (76,77). That conversation was the seed that led to many papers on the importance of the basement membrane throughout metazoan animals including this paper fifty-two years later.

## Data Availability

The shotgun transcriptome from wild type adult *Myxine glutinosa* Atlantic hagfish has been deposited at DDBJ/EMBL/GenBank under the accession GKQO00000000. The version described in this paper is the first version, GKQO01000000. The dogfish Transcriptome Shotgun Assembly project has been deposited at DDBJ/EMBL/GenBank under the accession GKOS00000000. The version described in this paper is the first version, GKOS01000000. Proteomics data for shark has been submitted to the ProteomeXchange, projects PXD0441912 and PXD042111.

## Author contributions

P.S.P., B.G.H., C.E.D., A.L.F., P.M., and J.K.H. conceptualization; P.S.P., C.E.D., A.L.F., and S.C. methodology; P.S.P., C.E.D., A.L.F.,S.C., J.G., Aspirnauts, and S.P.B. investigation; E. N. P., and S.C. data curation; P.S.P., E. N. P., and A.L.F. visualization; P.S.P., E. N. P., and B.G.H. writing-original draft; P.S.P., E. N. P., S. P. B., J.K.H., and B. G. H. writing-reviewing and editing; B.G.H., J. K. H. project administration; J. K. H., S. P. B., and B. G. H. funding acquisition.

## Funding and additional information

This work was supported by the National Institutes of Health grants R01DK018381 and R01DK131101 to B. G. H. and S. P. B. The Aspirnaut students were supported by NIH grant R25DK096999 to B. G. H. and by Lu-Springer Family Foundation.

## Notes

### Competing Interest Statement

The authors have declared no competing interest.

### Summary of Updates

The manuscript has been updated and edited prior to submission for publication.

## References

1. Fidler, A. L., Darris, C. E., Chetyrkin, S. V., Pedchenko, V. K., Boudko, S. P., Brown, K. L., Gray Jerome, W., Hudson, J. K., Rokas, A., and Hudson, B. G. (2017) Collagen IV and basement membrane at the evolutionary dawn of metazoan tissues. Elife 6

2. Fidler, A. L., Boudko, S. P., Rokas, A., and Hudson, B. G. (2018) The triple helix of collagens - an ancient protein structure that enabled animal multicellularity and tissue evolution. J Cell Sci 131

3. Kefalides, N. A. (1968) Isolation and characterization of the collagen from glomerular basement membrane. Biochemistry 7, 3103–3112

4. Kefalides, N. A. (1973) Structure and biosynthesis of basement membranes. International review of connective tissue research 6, 63–104

5. Spiro, R. G. (1967) Studies on the renal glomerular basement membrane. Nature of the carbohydrate units and their attachment to the peptide portion. The Journal of biological chemistry 242, 1923–1932

6. Beisswenger, P. G., and Spiro, R. G. (1970) Human glomerular basement membrane: chemical alteration in diabetes mellitus. Science 168, 596–598

7. Hudson, B. G., and Spiro, R. G. (1972) Studies on the native and reduced alkylated renal glomerular basement membrane. Solubility, subunit size, and reaction with cyanogen bromide. The Journal of biological chemistry 247, 4229–4238

8. Spiro, R. G. (1973) Biochemistry of the renal glomerular basement membrane and its alterations in diabetes mellitus. The New England journal of medicine 288, 1337–1342

9. Mott, J. D., Khalifah, R. G., Nagase, H., Shield, C. F., 3rd, Hudson, J. K., and Hudson, B. G. (1997) Nonenzymatic glycation of type IV collagen and matrix metalloproteinase susceptibility. Kidney Int 52, 1302–1312

10. Timpl, R., Wiedemann, H., van Delden, V., Furthmayr, H., and Kuhn, K. (1981) A network model for the organization of type IV collagen molecules in basement membranes. European journal of biochemistry 120, 203–211

11. Yurchenco, P. D., and Ruben, G. C. (1987) Basement membrane structure in situ: evidence for lateral associations in the type IV collagen network. The Journal of cell biology 105, 2559–2568

12. McCoy, R. C., Johnson, H. K., Stone, W. J., and Wilson, C. B. (1982) Absence of nephritogenic GBM antigen(s) in some patients with hereditary nephritis. Kidney Int 21, 642–652

13. Butkowski, R. J., Langeveld, J. P., Wieslander, J., Hamilton, J., and Hudson, B. G. (1987) Localization of the Goodpasture epitope to a novel chain of basement membrane collagen. The Journal of biological chemistry 262, 7874–7877

14. Saus, J., Wieslander, J., Langeveld, J. P., Quinones, S., and Hudson, B. G. (1988) Identification of the Goodpasture antigen as the alpha 3(IV) chain of collagen IV. The Journal of biological chemistry 263, 13374–13380

15. Barker, D. F., Hostikka, S. L., Zhou, J., Chow, L. T., Oliphant, A. R., Gerken, S. C., Gregory, M. C., Skolnick, M. H., Atkin, C. L., and Tryggvason, K. (1990) Identification of mutations in the COL4A5 collagen gene in Alport syndrome. Science 248, 1224–1227

16. Gunwar, S., Saus, J., Noelken, M. E., and Hudson, B. G. (1990) Glomerular basement membrane. Identification of a fourth chain, alpha 4, of type IV collagen. The Journal of biological chemistry 265, 5466–5469

17. Hostikka, S. L., Eddy, R. L., Byers, M. G., Hoyhtya, M., Shows, T. B., and Tryggvason, K. (1990) Identification of a distinct type IV collagen alpha chain with restricted kidney distribution and assignment of its gene to the locus of X chromosome-linked Alport syndrome. Proc Natl Acad Sci U S A 87, 1606–1610

18. Myers, J. C., Jones, T. A., Pohjolainen, E. R., Kadri, A. S., Goddard, A. D., Sheer, D., Solomon, E., and Pihlajaniemi, T. (1990) Molecular cloning of alpha 5(IV) collagen and assignment of the gene to the region of the X chromosome containing the Alport syndrome locus. Am J Hum Genet 46, 1024–1033

19. Hudson, B. G., Tryggvason, K., Sundaramoorthy, M., and Neilson, E. G. (2003) Alport’s syndrome, Goodpasture’s syndrome, and type IV collagen. The New England journal of medicine 348, 2543–2556

20. Hudson, B. G., Reeders, S. T., and Tryggvason, K. (1993) Type IV collagen: structure, gene organization, and role in human diseases. Molecular basis of Goodpasture and Alport syndromes and diffuse leiomyomatosis. The Journal of biological chemistry 268, 26033–26036

21. Poschl, E., Pollner, R., and Kuhn, K. (1988) The genes for the alpha 1(IV) and alpha 2(IV) chains of human basement membrane collagen type IV are arranged head-to-head and separated by a bidirectional promoter of unique structure. EMBO J 7, 2687–2695

22. Burbelo, P. D., Martin, G. R., and Yamada, Y. (1988) Alpha 1(IV) and alpha 2(IV) collagen genes are regulated by a bidirectional promoter and a shared enhancer. Proc Natl Acad Sci U S A 85, 9679–9682

23. Khoshnoodi, J., Cartailler, J. P., Alvares, K., Veis, A., and Hudson, B. G. (2006) Molecular recognition in the assembly of collagens: terminal noncollagenous domains are key recognition modules in the formation of triple helical protomers. The Journal of biological chemistry 281, 38117–38121

24. Khoshnoodi, J., Hill, S., Tryggvason, K., Hudson, B., and Friedman, D. B. (2007) Identification of N-linked glycosylation sites in human nephrin using mass spectrometry. J Mass Spectrom 42, 370–379

25. Pokidysheva, E. N., Redhair, N., Ailsworth, O., Page-McCaw, P., Rollins-Smith, L., Jamwal, V. S., Ohta, Y., Bachinger, H. P., Murawala, P., Flajnik, M., Fogo, A. B., Abrahamson, D., Hudson, J. K., Boudko, S. P., and Hudson, B. G. (2023) Collagen IV of basement membranes: II. Emergence of collagen IV(alpha345) enabled the assembly of a compact GBM as an ultrafilter in mammalian kidneys. The Journal of biological chemistry 299, 105459

26. Puapatanakul, P., and Miner, J. H. (2024) Alport syndrome and Alport kidney diseases - elucidating the disease spectrum. Curr Opin Nephrol Hypertens 33, 283–290

27. Brown, K. L., Cummings, C. F., Vanacore, R. M., and Hudson, B. G. (2017) Building collagen IV smart scaffolds on the outside of cells. Protein Sci 26, 2151–2161

28. Emsley, J., Knight, C. G., Farndale, R. W., Barnes, M. J., and Liddington, R. C. (2000) Structural basis of collagen recognition by integrin alpha2beta1. Cell 101, 47–56

29. McCall, A. S., Cummings, C. F., Bhave, G., Vanacore, R., Page-McCaw, A., and Hudson, B. G. (2014) Bromine is an essential trace element for assembly of collagen IV scaffolds in tissue development and architecture. Cell 157, 1380–1392

30. Poschl, E., Schlotzer-Schrehardt, U., Brachvogel, B., Saito, K., Ninomiya, Y., and Mayer, U. (2004) Collagen IV is essential for basement membrane stability but dispensable for initiation of its assembly during early development. Development 131, 1619–1628

31. Chen, Z., Migeon, T., Verpont, M. C., Zaidan, M., Sado, Y., Kerjaschki, D., Ronco, P., and Plaisier, E. (2016) HANAC Syndrome Col4a1 Mutation Causes Neonate Glomerular Hyperpermeability and Adult Glomerulocystic Kidney Disease. J Am Soc Nephrol 27, 1042–1054

32. Jeanne, M., and Gould, D. B. (2017) Genotype-phenotype correlations in pathology caused by collagen type IV alpha 1 and 2 mutations. Matrix Biol 57-58, 29-44

33. Plaisier, E., and Ronco, P. (1993) COL4A1-Related Disorders. in GeneReviews((R)) (Adam, M. P., Feldman, J., Mirzaa, G. M., Pagon, R. A., Wallace, S. E., and Amemiya, A. eds.), Seattle (WA). pp

34. Ninomiya, Y., Kagawa, M., Iyama, K., Naito, I., Kishiro, Y., Seyer, J. M., Sugimoto, M., Oohashi, T., and Sado, Y. (1995) Differential expression of two basement membrane collagen genes, COL4A6 and COL4A5, demonstrated by immunofluorescence staining using peptide-specific monoclonal antibodies. The Journal of cell biology 130, 1219–1229

35. Cui, Z., Zhao, M. H., Jia, X. Y., Wang, M., Hu, S. Y., Wang, S. X., Yu, F., Brown, K. L., Hudson, B. G., and Pedchenko, V. (2016) Antibodies to alpha5 chain of collagen IV are pathogenic in Goodpasture’s disease. J Autoimmun 70, 1–11

36. Gunwar, S., Bejarano, P. A., Kalluri, R., Langeveld, J. P., Wisdom, B. J., Jr., Noelken, M. E., and Hudson, B. G. (1991) Alveolar basement membrane: molecular properties of the noncollagenous domain (hexamer) of collagen IV and its reactivity with Goodpasture autoantibodies. Am J Respir Cell Mol Biol 5, 107–112

37. Hellmark, T., Burkhardt, H., and Wieslander, J. (1999) Goodpasture disease. Characterization of a single conformational epitope as the target of pathogenic autoantibodies. The Journal of biological chemistry 274, 25862–25868

38. Morrison, K. E., Mariyama, M., Yang-Feng, T. L., and Reeders, S. T. (1991) Sequence and localization of a partial cDNA encoding the human alpha 3 chain of type IV collagen. Am J Hum Genet 49, 545–554

39. Netzer, K. O., Leinonen, A., Boutaud, A., Borza, D. B., Todd, P., Gunwar, S., Langeveld, J. P., and Hudson, B. G. (1999) The goodpasture autoantigen. Mapping the major conformational epitope(s) of alpha3(IV) collagen to residues 17-31 and 127-141 of the NC1 domain. The Journal of biological chemistry 274, 11267–11274

40. Pedchenko, V., Bondar, O., Fogo, A. B., Vanacore, R., Voziyan, P., Kitching, A. R., Wieslander, J., Kashtan, C., Borza, D. B., Neilson, E. G., Wilson, C. B., and Hudson, B. G. (2010) Molecular architecture of the Goodpasture autoantigen in anti-GBM nephritis. The New England journal of medicine 363, 343–354

41. Turner, N., Mason, P. J., Brown, R., Fox, M., Povey, S., Rees, A., and Pusey, C. D. (1992) Molecular cloning of the human Goodpasture antigen demonstrates it to be the alpha 3 chain of type IV collagen. J Clin Invest 89, 592–601

42. Savige, J., Storey, H., Il Cheong, H., Gyung Kang, H., Park, E., Hilbert, P., Persikov, A., Torres-Fernandez, C., Ars, E., Torra, R., Hertz, J. M., Thomassen, M., Shagam, L., Wang, D., Wang, Y., Flinter, F., and Nagel, M. (2016) X-Linked and Autosomal Recessive Alport Syndrome: Pathogenic Variant Features and Further Genotype-Phenotype Correlations. PLoS One 11, e0161802

43. Savige, J., Gregory, M., Gross, O., Kashtan, C., Ding, J., and Flinter, F. (2013) Expert guidelines for the management of Alport syndrome and thin basement membrane nephropathy. J Am Soc Nephrol 24, 364–375

44. Savige, J. (2014) Alport syndrome: its effects on the glomerular filtration barrier and implications for future treatment. J Physiol 592, 4013–4023

45. Kashtan, C. E., Ding, J., Garosi, G., Heidet, L., Massella, L., Nakanishi, K., Nozu, K., Renieri, A., Rheault, M., Wang, F., and Gross, O. (2018) Alport syndrome: a unified classification of genetic disorders of collagen IV alpha345: a position paper of the Alport Syndrome Classification Working Group. Kidney Int 93, 1045–1051

46. Groopman, E. E., Marasa, M., Cameron-Christie, S., Petrovski, S., Aggarwal, V. S., Milo-Rasouly, H., Li, Y., Zhang, J., Nestor, J., Krithivasan, P., Lam, W. Y., Mitrotti, A., Piva, S., Kil, B. H., Chatterjee, D., Reingold, R., Bradbury, D., DiVecchia, M., Snyder, H., Mu, X., Mehl, K., Balderes, O., Fasel, D. A., Weng, C., Radhakrishnan, J., Canetta, P., Appel, G. B., Bomback, A. S., Ahn, W., Uy, N. S., Alam, S., Cohen, D. J., Crew, R. J., Dube, G. K., Rao, M. K., Kamalakaran, S., Copeland, B., Ren, Z., Bridgers, J., Malone, C. D., Mebane, C. M., Dagaonkar, N., Fellstrom, B. C., Haefliger, C., Mohan, S., Sanna-Cherchi, S., Kiryluk, K., Fleckner, J., March, R., Platt, A., Goldstein, D. B., and Gharavi, A. G. (2019) Diagnostic Utility of Exome Sequencing for Kidney Disease. The New England journal of medicine 380, 142–151

47. Pokidysheva, E. N., Seeger, H., Pedchenko, V., Chetyrkin, S., Bergmann, C., Abrahamson, D., Cui, Z. W., Delpire, E., Fervenza, F. C., Fidler, A. L., Fogo, A. B., Gaspert, A., Grohmann, M., Gross, O., Haddad, G., Harris, R. C., Kashtan, C., Kitching, A. R., Lorenzen, J. M., McAdoo, S., Pusey, C. D., Segelmark, M., Simmons, A., Voziyan, P. A., Wagner, T., Wuthrich, R. P., Zhao, M. H., Boudko, S. P., Kistler, A. D., and Hudson, B. G. (2021) Collagen IV(alpha345) dysfunction in glomerular basement membrane diseases. I. Discovery of a COL4A3 variant in familial Goodpasture’s and Alport diseases. The Journal of biological chemistry 296, 100590

48. Damas, J., Corbo, M., Kim, J., Turner-Maier, J., Farre, M., Larkin, D. M., Ryder, O. A., Steiner, C., Houck, M. L., Hall, S., Shiue, L., Thomas, S., Swale, T., Daly, M., Korlach, J., Uliano-Silva, M., Mazzoni, C. J., Birren, B. W., Genereux, D. P., Johnson, J., Lindblad-Toh, K., Karlsson, E. K., Nweeia, M. T., Johnson, R. N., Zoonomia, C., and Lewin, H. A. (2022) Evolution of the ancestral mammalian karyotype and syntenic regions. Proc Natl Acad Sci U S A 119, e2209139119

49. Simakov, O., Marletaz, F., Yue, J. X., O’Connell, B., Jenkins, J., Brandt, A., Calef, R., Tung, C. H., Huang, T. K., Schmutz, J., Satoh, N., Yu, J. K., Putnam, N. H., Green, R. E., and Rokhsar, D. S. (2020) Deeply conserved synteny resolves early events in vertebrate evolution. Nat Ecol Evol 4, 820–830

50. Chakraborty, M., Baldwin-Brown, J. G., Long, A. D., and Emerson, J. J. (2016) Contiguous and accurate de novo assembly of metazoan genomes with modest long read coverage. Nucleic Acids Res 44, e147

51. Zhou, J., Mochizuki, T., Smeets, H., Antignac, C., Laurila, P., de Paepe, A., Tryggvason, K., and Reeders, S. T. (1993) Deletion of the paired alpha 5(IV) and alpha 6(IV) collagen genes in inherited smooth muscle tumors. Science 261, 1167–1169

52. Mariyama, M., Zheng, K., Yang-Feng, T. L., and Reeders, S. T. (1992) Colocalization of the genes for the alpha 3(IV) and alpha 4(IV) chains of type IV collagen to chromosome 2 bands q35-q37. Genomics 13, 809–813

53. Al-Salam, A., and Irwin, D. M. (2017) Evolution of the vertebrate insulin receptor substrate (Irs) gene family. BMC Evol Biol 17, 148

54. Boudko, S. P., Danylevych, N., Hudson, B. G., and Pedchenko, V. K. (2018) Basement membrane collagen IV: Isolation of functional domains. Methods Cell Biol 143, 171–185

55. Fidler, A. L., Vanacore, R. M., Chetyrkin, S. V., Pedchenko, V. K., Bhave, G., Yin, V. P., Stothers, C. L., Rose, K. L., McDonald, W. H., Clark, T. A., Borza, D. B., Steele, R. E., Ivy, M. T., Aspirnauts, Hudson, J. K., and Hudson, B. G. (2014) A unique covalent bond in basement membrane is a primordial innovation for tissue evolution. Proc Natl Acad Sci U S A 111, 331–336

56. Gunwar, S., Ballester, F., Noelken, M. E., Sado, Y., Ninomiya, Y., and Hudson, B. G. (1998) Glomerular basement membrane. Identification of a novel disulfide-cross-linked network of alpha3, alpha4, and alpha5 chains of type IV collagen and its implications for the pathogenesis of Alport syndrome. The Journal of biological chemistry 273, 8767–8775

57. Kwong, R. W., Kumai, Y., and Perry, S. F. (2013) The role of aquaporin and tight junction proteins in the regulation of water movement in larval zebrafish (Danio rerio). PLoS One 8, e70764

58. Hillyard, S. M., Nadja & Tanaka, Shigeyasu & Larsen, Erik Hviid. (2008) Osmotic and Ion Regulation in Amphibians.

59. Zerbino, D. R., and Birney, E. (2008) Velvet: algorithms for de novo short read assembly using de Bruijn graphs. Genome Res 18, 821–829

60. Garber, M., Grabherr, M. G., Guttman, M., and Trapnell, C. (2011) Computational methods for transcriptome annotation and quantification using RNA-seq. Nat Methods 8, 469–477

61. Schulz, M. H., Zerbino, D. R., Vingron, M., and Birney, E. (2012) Oases: robust de novo RNA-seq assembly across the dynamic range of expression levels. Bioinformatics 28, 1086–1092

62. Altschul, S. F., Gish, W., Miller, W., Myers, E. W., and Lipman, D. J. (1990) Basic local alignment search tool. J Mol Biol 215, 403–410

63. O’Farrell, P. Z., Goodman, H. M., and O’Farrell, P. H. (1977) High resolution two-dimensional electrophoresis of basic as well as acidic proteins. Cell 12, 1133–1141

64. Timoneda, J., Gunwar, S., Monfort, G., Saus, J., Noelken, M. E., and Hudson, B. G. (1990) Unusual dissociative behavior of the noncollagenous domain (hexamer) of basement membrane collagen during electrophoresis and chromatofocusing. Connect Tissue Res 24, 169–186

65. Cox, J., and Mann, M. (2008) MaxQuant enables high peptide identification rates, individualized p.p.b.-range mass accuracies and proteome-wide protein quantification. Nat Biotechnol 26, 1367–1372

66. Cox, J., Hein, M. Y., Luber, C. A., Paron, I., Nagaraj, N., and Mann, M. (2014) Accurate proteome-wide label-free quantification by delayed normalization and maximal peptide ratio extraction, termed MaxLFQ. Mol Cell Proteomics 13, 2513–2526

67. Babicki, S., Arndt, D., Marcu, A., Liang, Y., Grant, J. R., Maciejewski, A., and Wishart, D. S. (2016) Heatmapper: web-enabled heat mapping for all. Nucleic Acids Res 44, W147–153

68. Martin, F. J., Amode, M. R., Aneja, A., Austine-Orimoloye, O., Azov, A. G., Barnes, I., Becker, A., Bennett, R., Berry, A., Bhai, J., Bhurji, S. K., Bignell, A., Boddu, S., Branco Lins, P. R., Brooks, L., Ramaraju, S. B., Charkhchi, M., Cockburn, A., Da Rin Fiorretto, L., Davidson, C., Dodiya, K., Donaldson, S., El Houdaigui, B., El Naboulsi, T., Fatima, R., Giron, C. G., Genez, T., Ghattaoraya, G. S., Martinez, J. G., Guijarro, C., Hardy, M., Hollis, Z., Hourlier, T., Hunt, T., Kay, M., Kaykala, V., Le, T., Lemos, D., Marques-Coelho, D., Marugan, J. C., Merino, G. A., Mirabueno, L. P., Mushtaq, A., Hossain, S. N., Ogeh, D. N., Sakthivel, M. P., Parker, A., Perry, M., Pilizota, I., Prosovetskaia, I., Perez-Silva, J. G., Salam, A. I. A., Saraiva-Agostinho, N., Schuilenburg, H., Sheppard, D., Sinha, S., Sipos, B., Stark, W., Steed, E., Sukumaran, R., Sumathipala, D., Suner, M. M., Surapaneni, L., Sutinen, K., Szpak, M., Tricomi, F. F., Urbina-Gomez, D., Veidenberg, A., Walsh, T. A., Walts, B., Wass, E., Willhoft, N., Allen, J., Alvarez-Jarreta, J., Chakiachvili, M., Flint, B., Giorgetti, S., Haggerty, L., Ilsley, G. R., Loveland, J. E., Moore, B., Mudge, J. M., Tate, J., Thybert, D., Trevanion, S. J., Winterbottom, A., Frankish, A., Hunt, S. E., Ruffier, M., Cunningham, F., Dyer, S., Finn, R. D., Howe, K. L., Harrison, P. W., Yates, A. D., and Flicek, P. (2023) Ensembl 2023. Nucleic Acids Res 51, D933–D941

69. Rangwala, S. H., Kuznetsov, A., Ananiev, V., Asztalos, A., Borodin, E., Evgeniev, V., Joukov, V., Lotov, V., Pannu, R., Rudnev, D., Shkeda, A., Weitz, E. M., and Schneider, V. A. (2021) Accessing NCBI data using the NCBI Sequence Viewer and Genome Data Viewer (GDV). Genome Res 31, 159–169

70. Altschul, S. F., and Lipman, D. J. (1990) Protein database searches for multiple alignments. Proc Natl Acad Sci U S A 87, 5509–5513

71. Hunter, J. D. (2007) Matplotlib: A 2D Graphics Environment. in Computing in Science & Engineering. pp

72. Lu, S., Wang, J., Chitsaz, F., Derbyshire, M. K., Geer, R. C., Gonzales, N. R., Gwadz, M., Hurwitz, D. I., Marchler, G. H., Song, J. S., Thanki, N., Yamashita, R. A., Yang, M., Zhang, D., Zheng, C., Lanczycki, C. J., and Marchler-Bauer, A. (2020) CDD/SPARCLE: the conserved domain database in 2020. Nucleic Acids Res 48, D265–D268

73. Federhen, S. (2012) The NCBI Taxonomy database. Nucleic Acids Res 40, D136–143

74. Schoch, C. L., Ciufo, S., Domrachev, M., Hotton, C. L., Kannan, S., Khovanskaya, R., Leipe, D., McVeigh, R., O’Neill, K., Robbertse, B., Sharma, S., Soussov, V., Sullivan, J. P., Sun, L., Turner, S., and Karsch-Mizrachi, I. (2020) NCBI Taxonomy: a comprehensive update on curation, resources and tools. Database (Oxford) 2020

75. Letunic, I., and Bork, P. (2021) Interactive Tree Of Life (iTOL) v5: an online tool for phylogenetic tree display and annotation. Nucleic Acids Res 49, W293–W296

76. Hung, C. H., Ohno, M., Freytag, J. W., and Hudson, B. G. (1977) Intestinal basement membrane of Ascaris suum. Analysis of polypeptide components. The Journal of biological chemistry 252, 3995–4001

77. Benigno D. Peczon, J. H. V., Calvin G. Beams, Jr. Billy G. Hudson. (1975) Intestinal basement membrane of Ascaris suum. Preparation, morphology, and composition. Biochemistry 14

78. Boudko, S. P., Bauer, R., Chetyrkin, S. V., Ivanov, S., Smith, J., Voziyan, P. A., and Hudson, B. G. (2021) Collagen IV(alpha345) dysfunction in glomerular basement membrane diseases. II. Crystal structure of the alpha345 hexamer. The Journal of biological chemistry 296, 100591

79. Sundaramoorthy, M., Meiyappan, M., Todd, P., and Hudson, B. G. (2002) Crystal structure of NC1 domains. Structural basis for type IV collagen assembly in basement membranes. The Journal of biological chemistry 277, 31142–31153

80. Hermetz, K. E., Newman, S., Conneely, K. N., Martin, C. L., Ballif, B. C., Shaffer, L. G., Cody, J. D., and Rudd, M. K. (2014) Large inverted duplications in the human genome form via a fold-back mechanism. PLoS Genet 10, e1004139

81. Ballif, B. C., Yu, W., Shaw, C. A., Kashork, C. D., and Shaffer, L. G. (2003) Monosomy 1p36 breakpoint junctions suggest pre-meiotic breakage-fusion-bridge cycles are involved in generating terminal deletions. Hum Mol Genet 12, 2153–2165

