## Supplementary Figures for "Collagen IV of basement membranes: I. Origin and diversification of COL4A genes enabling animal evolution and adaptation"

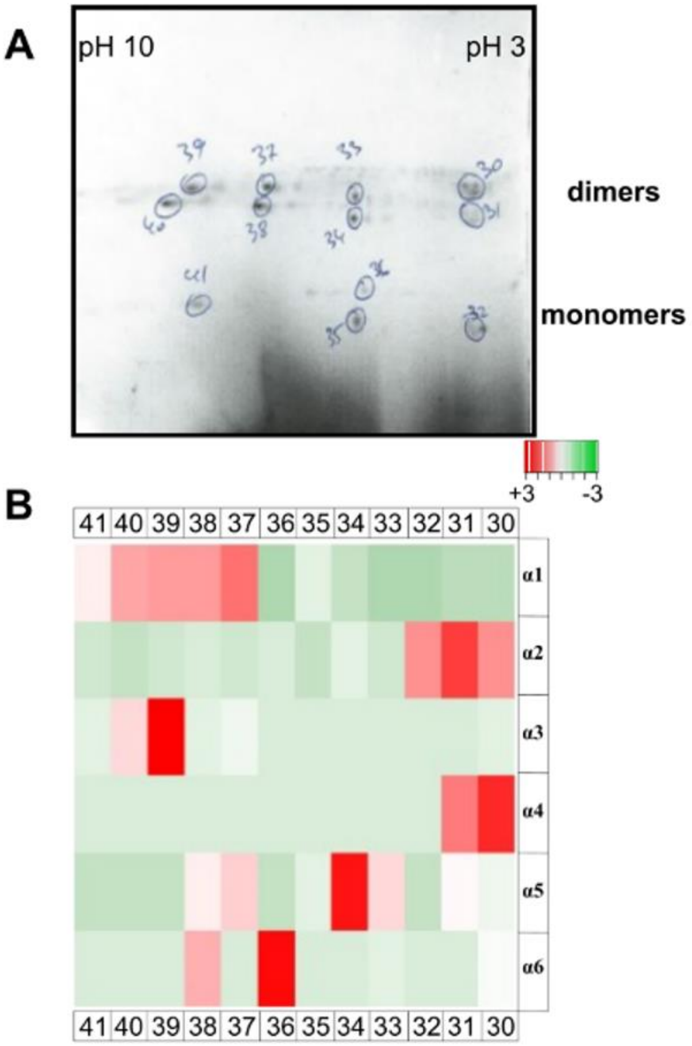


**Supplementary Figure 1.**

A. Colloidal Coomassie stained 2D-NEPHGE gel with spots that were excised for mass spectroscopy (indicated with the numbered circles).
B. A heat map (heatmapper. ca) built from label free quantification (LFQ) of protein level data. A custom dogfish protein database was used to define chain composition in each 2D spot.

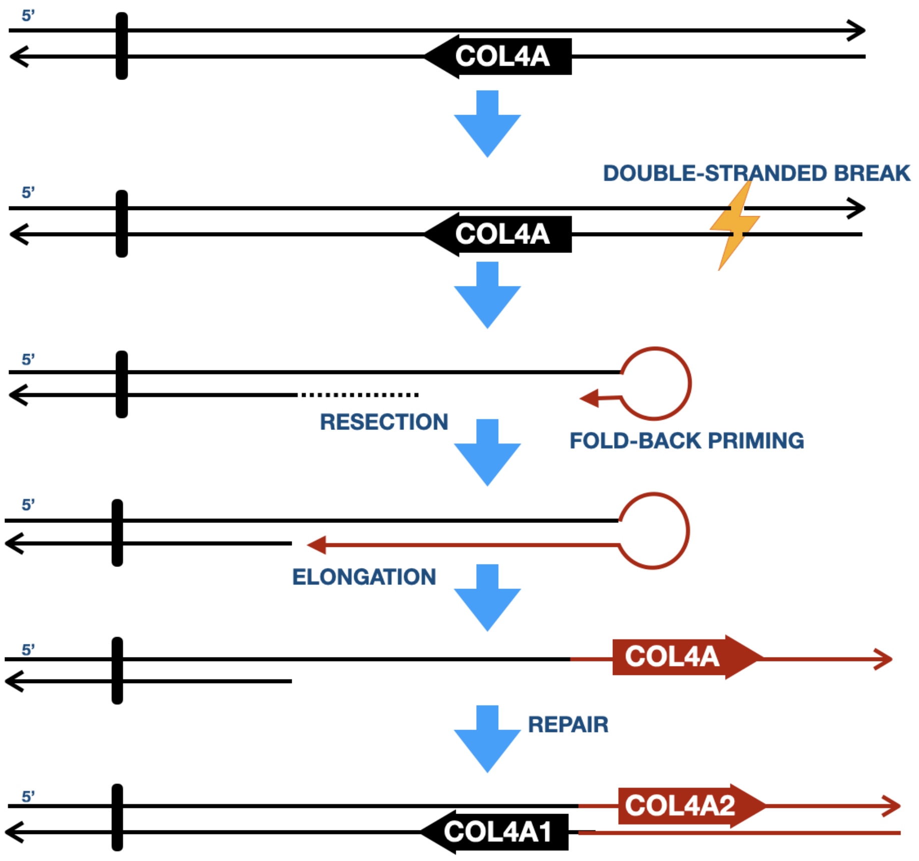


**Supplementary Figure 2. Hypothesized mechanisms for COL4A gene duplication.**

Proposed mechanisms for the generation of the COL4A gene pair in the head-to-head gene arrangement. (A) The original COL4A gene is found likely in a telomeric position on the chromosome and its gene duplication is initiated through a double stranded break telomeric to the ancestral COL4A gene. Repair of the double stranded break proceeds through fold-back priming from the 3' end of the top strand, resection and elongation through the COL4A gene (red strand), followed by resolution through multiple possible mechanisms (80). A similar model could occur in meiosis through creation of a dicentric chromatid that breaks during anaphase results in one gamete carrying a deletion for the ancestral COL4A gene and one carrying the duplicated COL4A⟨1|2⟩ gene pair (81).

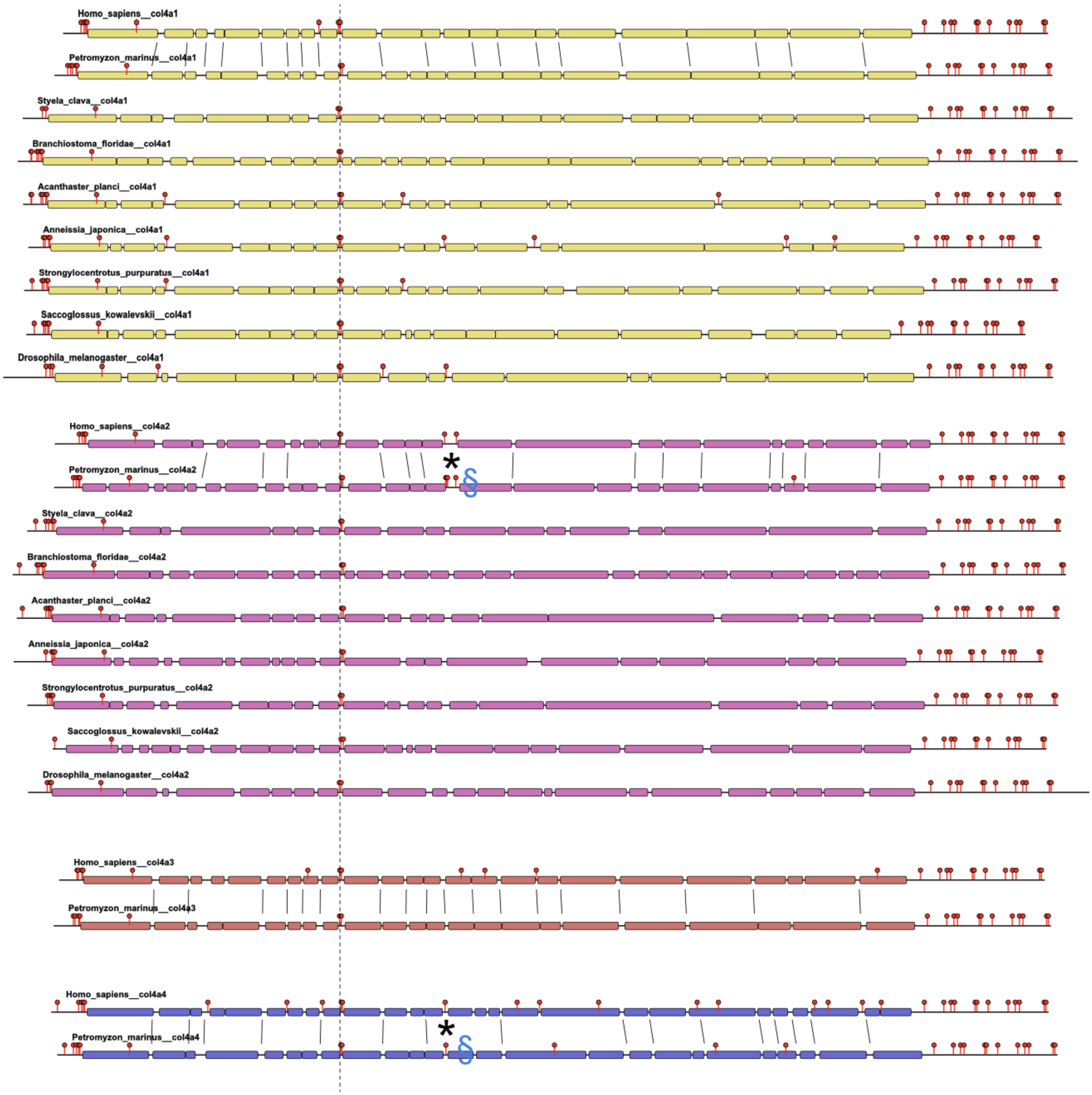


**Supplementary Figure 3. Comparison COL4A⟨1|2⟩ domain structure within Deuterostomes.**

Predicted collagen domains are shown as blocks with interruptions as lines, the collagens are aligned to the highly conserved cysteine pair. Conservation in placement of the cysteine residues can be seen in the presence of the red flags. The 7S and NC1 domains are found at the N and C termini. Block diagram showing the domain structure of the predicted products of the COL4A⟨1|2⟩ loci in basal Deuterostomes. Collagen helical domains are indicated as blocks (Yellow COL4A1, pink COL4A2).

Alignment of the gaps is indicated between the human and cyclostome sequences and the presence of the cys-loop-cys motif is indicated by *. Position of Cys residues and interruptions to the collagen blocks is observed between the human and cyclostome COL4A1 and COL4A2 genes, consistent with a correct assignment of the genes to their respective COL4A gene families.
