## Supplemental Text for "Collagen IV of basement membranes: I. Origin and diversification of COL4A genes enabling animal evolution and adaptation"

### Supplementary Text

Peptide sequences of Dogfish and Hagfish COL4A genes determined in this study.

```
>COL4A1-like_[Squalus acanthias]
MEAAACHGCTSSSKCDCSGVKGEKGAAGLPGFAGPLGMPGFPGVEGPVGSVGPKGDSGEPGLVGKKGIRGPAGSPGFPGT
PGLPGLPGQDGPSGPGGIPGCNGTKGEKGPLGPPGFPGLQGPSGQPGIPGMKGDPGEVIDISVSKGERGFPGPPGKVGTP
GISGPIGPPGPSGYPGPSGPPGIPGLPGPKGNMGVQFYGPKGQKGEQGVRGPPGAPGPGEHTSSPETKVQKGEKGQKGDH
GLPGAKGFPGSPGHSSIGMNGDKGELGEPGKRGKPGKDGDPGFPGRDGQSGYPGEPGFIGQDGPKGEKGNRGIDGPPGVI
IDSDGQAIYGTQGEPGSPGPPGLKGEQGPSGPPGLAGVAGPPGLSQQGEKGLPGFTGERGQKGEKGVPGLSLSGSPGRDG
QSGLPGPPGPPGPPVSAADCNSIPGPPGAAGPRGFQGETGLKGAKGDYCFHCNFTLSPGPAGPQGPPGRSGDPGIPGTKG
DHGPPGRVGFPGQTGATGVPGPMGIPGPKGDPGDIQTLPGLKGDRGDSGSPGTPGVPGLDGLPGKDGSSGFPGPKGESAT
IGFKGDPGIPGTPGLHGAPGDQGPPGLPGVGEPGFPGEKGSQGFPGRPGIPGTTGQKGESGNGISRPGSPGLPGTQGLRG
EPGGRGDPGLPGAPGVQGAVGPKGDPGPTGIGFPGPTGVKGVQGIPGQSGPPGLSGQPGYDGLPGQPGTPGIKGQAGVGL
PGPQGLPGTPGFDGIQGAKGDPGLPGIPGGPGRPGLPGFPGEKGSHGLVGVQGPVGPPGPPGIGAPGQVGNPGPQGPSGL
PGFAGPPGAKGDPGYPGLDIPGPQGEQGRQGSKGHPGPPGYPGPIGQPGRDGPSGLQGPKGEMGVMGTPGSQGFPGSPGS
AGFSGPKGDQGLAGLPGQSGREGEKGIKGETGLAGPPGVMDKSAMEKMKGQKGDIGDHGKPGASGTKGYQGIHGDPGMPG
KDGEPGLPGQAGPVGDSGNPGKPGSTGVPGDKGSIGEMGLPGISGPRGPPGDPGLEGIPGFTGLTGEKGDKGNPGIPGFG
VPGDPGPKGDSGLPGFPGDPGLTGDSGLPGLTGSLGAPGLKGGAGVPGFPGNPGLPGKKGGFGPVGEQGFSGLPGQPGPP
GPKGTVGVHGEKGSRGLDGIPGTIGEKGNQGLAGRDMPGAPGLPGTKGVKGDLGLPGYPGSPGVPGIKGERGFPGNEGHQ
GNPGLPGPPNVALEGPKGDKGTAGGTGRPGAPGVHGGPGAPGLDGTKGDRGEQGSSGFQGFPGPKGDPGVSGSPGSAGQP
GAPGLKGESGPHGVPGFIGMKGTSGPPGFPGIQGEGGPPGQLGPPGPVGPQGPLQIVKGDPGLPGRPGEDGAKGYPGFPG
VKGDVGLLGPIGDPGEEGAPGFDGRPGLKGELGPSGPTGSRGYPGPAGPEGAPGPRGLPGPASIAHGFLVTRHSQTTEVP
TCPHGTSKIYDGFSLLYVQGNERAHGQDLGTAGSCLRRFSTMPFLFCNINNVCNFASRNDYSYWLATPEPMPMSMAPVIG
QSIKPFISRCSVCEAPAMVIAVHSQTIQIPLCPIGWHSLWIGYSFMMHTSSGAEGSGQALSSPGSCLESFRSAPFIECHG
RGTCNYYANSYSFWLATVESSEMFRKPESETLKAGELRTRISRCQVCMKKT
>COL4A2-like_[Squalus acanthias]
MWCLLLQLLIPVLLADKVLAGGKSYSGPCNGRDCSAGCQCFPEKGARGRPGIPGVQGPQGPPGPRGRQGVPGYKGHKGYR
GTDGLTGHTGDKGPMGVPGFAGNDGIPGHPGQPGPRGPPGHDGCNGTRGVPGEMGPPGFPALLGFPGAPGRKGQKGEPFL
VTVERTRFRGDPGHAGAPGYPGTSGLPGQMGSRGPPGVLGPQGRPGPPGAQGPKGNDAVSFQGRTGDKGEAGLPGVPGPG
AGTGPPVTIYAKGEAGEKGERGNEGFEGEVGHPGKPGVPGPQGEKGEEGIMGLPGSRGSSGLDGPPGSVGATGLPGIRGL
PGVDGRLGMKGDRGDQGLPGRSITYSGSVPKGPKGFRGLEGDRGHQGDYGPHGPRGHPGPPGKTPSYRGFRGPQGSPGVT
GPKGYKGEVGLPALQPGFIGFPGISGPPGPPGLPGPASSAEFIGDPGRRGQRGYPGQRGGTGGKGEAGHCACIMEDAAPA
GPPGPVGVPGYPGRYGVTGQLGDEGAPGPFGVPGTPGAAGAVGARGPKGQKGDSIIITTITTKGSKGDRGAPGRTGAHGQ
PGLPGQDGKLGKQGFPGPSGDAQKGYPGDDGIRGGPGRKGYSGRPGLPGVGIRGPQGRLGLLGDLGLDGVPGIPGIPGLP
GTPGDCIDEQEILTREEDPSIFQSAIPCGTPGSPGQRGLPGLPGDKGVKGQLGFPGLSRPGFPGRKGFIGAPGPGTPGPP
GFVGPRGDQGLPGPPGQDGDAGLQGKPGRPGPLGAKGLSGEAIGALPGYQGESGHPGRPGAKGSQGDAGETGYRGADGML
GMPSPKGERGQPGSPGIAGLPGLKGQPGGLGQPGFPGLPGIPGFPGSVGSQGYIGEHGDRGPPGPAGMKGLPGIPGAKGL
LGDVGFPGPFAPVGMDGLPGLDGLKGLKGSPGMAGFPGARGVKGWHGIRGLKGDSGFPGSTGSSGPLGRGGLKGDRGANG
PPGPQPPIEPAMMERVKGDKGALGYSGESGFVGPRGSKGMPGLPGQLGHHGFPGVPGHQKGLPGFPGLPGETGSKGMPGS
TGPGMYGEKGVSGNPGLPGALGLPGPSGQKGTLGETMNMPGQSGRKGETGQLGSKGNAGLPGLPGKLGHPGEPGIEGMKG
TPGDPGLPGQQGSPGFPGTPGGRGSRGPDGSSGLSGYTGFPGYRGEKGWQGSEGLPGLDGLTGNKGFPGSPGVSGRGAQG
HLGESGEKGEQGVSSTQPGNPGLPGDAGPRGESGEPGPIGPTGPTGAPGPLKPSDQPGKPGDIGPRGADGPQGFPGQPGP
PGRWAPPGVKGEPGDQGRAGFPGPPGLSGDEGSPGLSGIDGISGNKGQRGDQGMMGYPGRKGPHGAQGAPGVKGPVGLAG
ILGLPGPRGPAPPPQRFPAPEGSRGLPGSPGRHGFPGDPGPKGNPGDLGPKGPKGRPGIPGRSNSPGTMGPRGEPGNIGQ
QGPLGFHGAPGTTGIRGMAGMPGRSVSVGYLLVKHSQSQQVPPCPLGMNRLWDGYSLLFVEGQEKSHNQDLGLAGSCMPR
FSTIPFVYCNINEVCYFASRNDKSYWLSTNAPIPMMPVGDEDIKPYISRCSVCEAPSLAVAVHSQDITIPMCPAGWRSLW
IGYSFLMHTAAGSEGGGQSLVSPGSCLEDFRTTPFVECNGARGTCHYFANKYSFWLSTIDPSQQFEELPVPETLKAGQLL
TRIGRCQVCMKNL
>COL4A3-like_[Squalus acanthias]
MPVPVPTLSCVLVILSIATLAEAACTCSGKSKCHCDGVKGNKGESGFPGLQGPPGIPGFPGPEGPPDQEGPKGNHGFPGP
PGMKGLRGPPGLPGYPGNPAVPGIPGVDGPLGPKGIPGCNGTKGDPGFPGIPGSEGLTGLQGHPGLAGQKGEPAPMVLLC
PGEKGNRGLQGHRGSPGRPGFPGLQGPAGPQGPLGPSGQPGRPGPKGNMGLFFVGFKGDKGVKGLPGPVYRQDRALTITE
KGQKGSRGPSGDPGFTGPPGLPGPVALGWKGDKGERGDSGALGKPGKSGTTGLPGFQGLKGDPGFPGFPGRDGRPGRKGV
IGRQGPPGPQGEEDNRDCRKGDRGCHGPTGPRGQPGLPGAVGQRGLPGKPGSSRRGPSGQPGYPGQQGPKGAKGDSGQIS
EGPQGPPGFPGLPGLPGPLGPPGENIRINGQPGPAGPPGPTGFVGFHGEGGYKGDKGELCITCNGISVPGPPGPPGPPGI
PGRPGFPGLKGNPGFTGEHGRTGFQGLPGYPGVGGSPGPKGDSAICGSQGETGNKGDPGFHGNTGKQGNDGIPGQPGPQG
PRGIKGDPTPFSPKGSMGPRGNSGNPGVPGPMGPSGPPGFGPPGAQGPKGNRGVLGFRGVDGVPGTKGSKGNGNVIPGPP
GFRGEMGGQGRSGIQGNPGPPGRDGSPGAPGEKGEIGPSEKGPPGPPGPKGNPGLSGSLGPPGTPGRSGAPGTPGLPGVT
GPKGVRGYGQPGNRGPPGPPGNVGLPGEKGDSGSCGLPGGEGRPGRTGPQGPKGELGSYGNQGATGPRGPRGRGIYGPAG
VEGPPGAPGLQGMTSRGPTGDPGIPGLDIPGPPGIPGLIGNPGCSGPPGPRGSPGHPGRPGTSGRSGPKGEPGVMGEPGT
PGLTGAQGHQGPRGDKGSEGRTGPKGEQGLLGMTGQRGEKGMKGVPCNMTVKGALGEPGLPGAKGRTGPKGETGACADPG
YIGKEGTPGPEGPAGPPGDPGITGNPGYRGSPGVTGSMGSMGLPGDKGDKGFPGTPGSLALPGPGGSSGPKGKKGEPSRT
DYGNTGPPGQKGNRGISSPKGSPGTPGSPGKPGLVGRSGAPGEKGDLGQSGCAGSSGPQGYKGLDGKQGLPGGPGVPGTP
GITGRIGPPGAMGAKGNQGSDGFPGLHGLKGDPGFPGSGYKGQKGRPGPPGLKGDIGLPGDDGGQGSPGPGGDPGPHGPP
GSCGEPGPPGPAGMSIQGPRGNRGSQGTPGSAGYRGPKGLLGQPGSDCIKGVKGSPGLPGVPGNPGDRGDPGAKGQLGIP
GTSGQGPKGERGSTGTPGFPGIQGDKGVQGQTGGAGQQGLLGKPGPKGLPSIVEIIRVKGDLGFPGPPGSEGQSGTPGSA
GPPGCIGNPGPPGPLGRNGPPGGVGQSGSKGSSGFLGEQGPKGYPGSPGNQGDWGHPGPSRPDIQGFIFTRHSQTIEIPL
CPGSTKQLYVGYSLLFLQGNKRAYGQDLGAAGSCLPRFSVMPFLFCDVNDVCNYASRNDYSYWLSTSKQMPMDMAPISGQ
ELQPYISRCIVCESPAMVFAVHSQTVQIPPCPRGWKSLWIGYSFVMHRSSGAEGSGQPLASPGSCLEEFRVAPFIECHGR
GTCNYYVNAYSFWLATLDPSQMFRKPRPQTLKAGELRSIISRCQVCTK
>COL4A4-like_[Squalus acanthias]
MALTGFLNRCLVQIAPLGPLWLLLLVQLVFLRGIHAGKIGYTGPCDGRDCSVCQCFPAKGSQGQPGLLGRQGPPGSPGHQ
GPHGPSGLKGQRGDRGLPGSAGIKGDGGPTGVPGFQGLDGIPGFPGLQGPRGHPGLDGCNGSRGDPGLPGIDRYPGRYGL
PGLPGPKGPKGDSITVTSSEGCLPGDQGFQGQMGFPGLPGNPGSPGIAGPRGQPGFPGRPGLQGLPGEQGIVFPGQKGEK
GEQGDPGHVELIQPPEEIVFKGDQGEKGPRGPPGPIGFPGPSDLPSYPGEKGEKGIIGFPGNRGDPGKEGLSGVPGRKGQ
HGPIGRPGSDGYPGTEGDPGNPGPAGPPGLTHLPCNPQKGDSGRRGFPGRNGEPGLPGPPGQPGRPGDSFGPGLPGPAGF
```

PGLDGPQGEKGRKGSDDCPKPDIIEGPRGRPGLGGPRGPHGSKGNPGFICETPGPPGPPGLKGNQGPPGQKGTDGLKGDL
GDCACHVDLRGTPGNSGTPGIPGNPGSPGRKGEQGDRGLPGSVGLPGLAGPPGRQGSLGAQGQKGEPSQDAKKGEKGIPG
DPGPAGARGRPGEPGQHGNPGLIGQRGVPGEAKKGFPGDKGLPGPPGPTAFPGPPGPSGVGYPGPKGQNGPVGGEGFPGP
KGAIGQRGQKGDSLCPSSPGPEGPKGMPGPPGPIGSPGILGRPGQIGFDGFKGLKGEPGLMAYPGPSGFQGSVGAPGCQG
PSGTSEDGAQGLPGLPGTPGERGPPGDSSDGPPGAPGQIGRRGSTGARGDTGDIVSPGSRGRDGVPGFPGLKGSQGDRGP
TGLPGNSGPPGFPADKGDPGPVGPPGRPGLRGPQGLSQKGFLGNPGPLGPAGLKGLQGIPGNPGTRGPIGIQGQPGPVGT
SGTPGFPGPAGSEGSQGIQGFPGASGPRGYPGIPGQQGLRGDDGRKGPDGKRGIYGNKGERGDQGDESMPMIFYVKGTSG
IPGLPGEHGFTGPRGEKGLPGNPGIPGFSGRPGLPGSKKGFPGMSGPSGPPGRQGFMGQKGQRGIIGFPGMNGEKGDIGQ
LGSPGLQGAPGCKGTKGDSGDSISFPGVPGPKGQIGDPGEPGVTGLPGQPGPSGIDGRSGAPGVKGIHGDPGMPGFPGIQ
GPPGDSGRLGPLGSPGNPGQPGSVRPPGPPGSKGQPGSPGLNGLNGLQGQKGTHGPPGPSEPGPSGSPGSSGSKGVKGAP
GLPGSSIPCPVGSPGPRGPEGSPGVRGAPGPPGLRGPPCSIISPGPPGSEGPPGFDGPPGPPGFPGDPGIPSIPFQGDPG
VPGTPGRSGAQGPPGEQGSCGFNGPPGPQGRKGEMGDQGYFGFPGLVGPLGAPGDRGGCGYPGPEGPQGDPGSPGSIISD
VGDVDMSHYKGDPGSPGLPGPPGPVETMGYSGPRGPPGLKGLKGEPGRAGADGAPGPPGPLGNLGKPGPRGITGDQGPPG
QQGPSGNPGTCLPFSAGFLIVVHSQSQKIPTCPQNMLSLWQGYSLLYLEGQEKSHNQDIGLAGSCLPMFSTMPFAYCSID
EVCHYATRNDKSYWLSTTAPIPMMPLIEHEIEPYISRCSVCEAPSVAVAVHSQDQTIPPCPSGWISLWAGYSFLMHTGAG
AEGGGQSLTSPGSCLEDFRSTPFIECQGARGTCHYFTDKYSFWLTTVQPNQQFEFSAPSETLKIEQLQRQRVSRCQVCLK
NK
>COL4A5-like_[Squalus acanthias]
MISPKSRAGKMKFEKSFTFGQLAARVFLLLSVLTVWVQNSEAAACHGCSTGSDCDCSGAKGEKGMRGLPGLEGHTGLPGF
PGPEGPPGSRGIKGDTGPPGFSGPKGIRGPPGLPGFPGTPGVPGLPGQDGPSGPPGIPGCNGTKGEQGFSGGPGFPGLQG
PPGPPGLPGYKGDPGEVLSSQLSGHKGDPGLPGLPGLPGPQGTPGSSGPLGPSGPSGQPGPPGSPGQPGQKGNMGLNFQG
PKGEKGDPGLQGPPGPPGQLEQLNSPGVDFQKGDKGDYGPPGSPGERGFPGPPGPPGFSGKGAKGEPGDAGKRGKPGKDG
ELGSPGFDGLPGTSGQPGGPGRDGAKGLKGEYGLPGPPGITFPRPGVTVGQKGDAGFPGSPGQSGERGPAGFPGLPGPPG
APGQPSVGSPGTPGFPGERGQKGDEGVPGYSLPGPAGPGGRPGPPGPPGPPGHSGPSVPGHGSQCQPGQPGLPGPRGSPG
FPGESGLKGDRGDTCINCLDTGDRFPGVPGPPGPHGIPGQPGLPGMKGDQGLPGPIGNLGLPGSPGRPGNPGSSGLKGDR
GDGFGYPGVKGDRGGSGFPGPPGLPGLDGTPGRDGVPGSPGQKGVPGGLAFKGGRGLTGDPGPLGFPGERGPTGPPGFGP
PGYPGDKGVQGVSGRPGAPGAPGIKGESGSTISEPGPAGPPGPHGESGLPGRPGDSGQPGQPGLSGLPGSKGDMGVPGIG
FPGPTGLKGLPGSPGLPGAPGNSGRPGRDGSPGQPGRSGSKGDPGFGLPGPSGPSGLRGNDGLPGPKGDSGFPGLPGQPG
RPGLDGTLGTKGDPGFGGPPGSTGPPGPPGIGGRGQPGPQGPPGLPGQPGRTGLPGPYGDKGDPGPPGLDIPGPPGDKGN
PGLPGSPGSNGLPGSPGRPARDGLPGPAGPKGDMGVMGTPGPSGPPGSSGVPGFQGAKGNEGFPGSPGNPGGPGVRGLKG
DAGLPGSPGTIDPLQYVAGQKGDPGFSGSPGLPGAKGFSGVPGNPGAPGQDGLPGLPGGLGPKGDTGFSGQPGSPGRPGP
KGSIGGMGLPGPPGNKGTPGTSGRPGQPGSPGRPGFQGEKGDTGVPVLGFPGSPGYKGDPGPPGFSGVPGSKGTPGSPGL
PGLPGAPGGKGDPGQPGFPGTPGIPGPKGIDGFPGSPGLIGPPGPPGEPSRPGSPGLPGEKGQPGRDGIPGPAGLKGDSG
PPGLGRPGSSGLLGIPGPKGDSGVPGIPGGPGAPGLKGDPGFPGFAGQQGPPGPPGPPGHSLEGPKGTPGSSGQPGRPGP
QGPIGQRGPPGFGGIKGEKGNSGLPGSPGFPGPKGEFGPPGIPGAPGQPGFNGPKGDPGVPGLPGFPGMKGSPGPFGLKG
TPGDLGPPGPPGQPGTPAPQVPLRSFKGERGFPGQPGQPGLKGLPGSPGPPGPTGPVGSSGDPGQEGLPGFSGPKGQKGD
SGPSGNPGQRGFPGPPGSDGTPGIPGPPGSASVAHGFLITRHSQSTEVPSCPSGTGVIYDGFSLLYVQGNERAHGQDLGT
AGSCLRRFSTMPFMFCNINNVCNFASRNDYSYWLSTPQPMPMSMSPVNGENIRPFISRCTVCEAPAMVIAVHSQTIQIPL
CPEGWASLWIGYSFMMHTSAGAEGSGQALASPGSCLEEFRSAPFIECHGRGTCNYYANSYSFWLATVEMAEMFSKPQSET
LKAGELRTRVSRCQVCMKRT
>COL4A6-like_[Squalus acanthias]
MKVPRKETVNKTKPINMSNGILLMVVACLAADLVQAGVSNVYFGPCEGRDCSAGCKCFPEKGSRGQPGLIGPQGRSGPPG
FSGPEGLSGPKGDKGNQGNPGVGGMKGEKGPIGVPGFVGMNGIPGHPGQQGPRGRAGLDGCNGTKGDSGFPGGAGYPGSL
GFPGEVGHKGAKGEPAYVSGGYGGLRGEPGLPGLDGFSGQKGYPGSYGPTGPRGRPGTTGRPGPPGSRGLKGNMGLGFQG
QKGGKGDVGLPGPPGRPGTIPHGSGPGINISIIGEKGDKGLPGAPGRPGIRGPPGYSDVNRAKGEKGIPGLPGPRGFSGL
DGIPGNPGRKGQAGFVGPNGRDGYPGLKGDHGEMGPPGPPAYVDGSGTILKGPRGDPSPPGLPGSPGSRGSRGQLGLPGP
PGFPARTQGSKGFSGFPGVGGPKGEKGMPGKTLFSQTGPQGQPGLPGPMGPPGAPAYSGPTDNRGEITGPVGPPGFPGPN
GPPGFRGSKGYKGEPGACACNGVIIGSSSQGPPGPPGAPGEVGFNSVKGQCGDPGPPGPPGQSGLPGLPGVPGGSGVKGQ
KGDSSGTSAKGPKGDPGTPGSRGPSGTPGKPGRDGFPGLPGPPGIPGDSGTGFPGVKGLPGSPGRKGPPGERGAPGIGLP
GPPGFQGPPGDPGFAGIPGPPGFRGLPGDCCCGETAREGDVHTEGGITLPCVIPGGRGLVGRPGSPGVPGSKGRPGFPGN
QGRPGFDGPKGPPGTPGLLGESGRPGFPGARGDQGLPGLPGLDGNEGPPGKMGAPGFSGQKGLPGDLYGAESGAPGQPGQ
PGKPGLKGAPGDSGLPGPRGFDGRPGFPNPKGERGVPGYPGRRGLPGPPGVSGPDVITGFPGEAGRPGEPGQPGFPGPKG
FVGDRGPSGPSGMKGFPGTPGNPGLLGRQGYPGAPGEPGREGVPNRYGLKGERGSPGVIGFPGMPGLPGRPGYDGRKGEL
GSPGRPGFPGSPGRSGEKGDRGSNGLPGPISVIENGRRRPGPRGVSGTPGVPGFSGPRGGKGFPGPPGSPGLPGLRGLPG
REKGRPGDFGIPGSPGSPGQFGLKGSPGIFGFPGMMGEKGDDGLPGSPGYPGELGLVGPKGERGDVLSVPGSLGPKGESG
QPGYPGTRGRDGLRGDPGFSGSPGTSGFKGSPGEGGRPGTPGGRGLQGTPGSPGYQGSRGPPGLPGRSIQYTPPGQKGLP
GSPGLDGLNGLPGPKGPPGTPGAGLPGQPGSPGLPGLKGEYGTPGTGIGAPGYPGEKGLRGDPGHGGSLGLPGKPGAPGI
TTASNIPGRPGDLGPPGYDGEAGLPGPTGSPGPPGLGFERGDPGNPGLPGVPGPQSIRGNIGPPGISGVPGPPGLKGQRA
EPGRMGFPGTDGPPGDQGYPGHKGDPGARGYPGNPGFPGQPADIEEEPPIPGASGLPGYDGESGPQGDPGSLGYLGFPGI
KGQPGIPGRVGPNGYPGPPGSLGDPGPDGYPGAPGLEGYPGKPGSAGLPGMSARSVNVGYTLVKHSQSDNVPPCPLGMNR
LWDGYSLLFVEGQEKAHNQDLGMAGSCLPRFTTMPFLYCNIDEVCHYANRNDKSYWLSTTAPIPMMPVSNLQIRQYISRC
SVCEVPSQAIAVHSQDITIPQCPEGWRSLWIGYSFLMHTAAGAEGGGQSLVSPGSCLEDFRATPFIECNGARGTCHYFAN
KYSFWLTIVEASRQFVELPPSETLKAGQLRTRVSRCQVCMKYL
>Col4aA_[Eptatretus burgeri]
MLRCTQLSGARSSRTVGVRGSADVLRLLVIVVALLGDTVTAGNDWNTGPCGGRDCSAGCECFPEKGSRGRPGSIGSRGRK
GLLGPPGMYGPTGVKGHRGSMGRPGHHGGKGDSGPIGVPGFPGIDGIPGHPGQSGPRGPAGADSCNGTQGEPGDPGYGSG
LPGLRGLPGLIGPKGQKGDPSYGDFTGVGNPGDPGIPGAPGLPGTPGSSGYEGPQGPMGLPGPPGYPGPIGLPGDTGFED
VSLQGQEGDKGDPGEQGPPGNSTYVPVSGPGIIELRGKKGHKGSQGNSGYFGAKGYRGTPGFTDGIGMNGEKGIIGLPGP
RGPPGADGIAVPGRKGATGYDGRMGPNGDRGMKGERGDRGLPGPITYLPPGIPQIKGYPGDPGPLGRPGLQGDPGFEGPR
GPPGPPGYRTGDRTDNDCAGQRGPPGPPGVFGPKGEMGDPGRPGVGQPGELGHDGEPGFQGSPGSPGPPGVVIPVRSNRT
GLPGISGQPGDIGSSGNHGSMGQKGDPGDCLCQTGPRPDVDDGFPGPPGPRGQQGNSGFPGLKGLPGDRGFPGMPGSRGY
DTGLPGEVGLKGPKGSKGESYFATEKGCKGKAGTPGNQGPSGPMGQPGRDGVPGFPGLKGPPGDGGVGYRGDKGFPGLQG
SPGRQGPDGAPGRGFPGRPGQRGPPGESGFIGMPGTPGPKGIPGDECAKKRHGLPGPPGPVGYTGPKGLPGIPGNDGQPG
FPGSKGERGDPGITDAPSLPGFPGIIGDPGGPGSRGSDGIPGGPGRPGNPGIPGAKGDPGDVLFSSTGPPGEQGRPGFPG
GKGEPGVSGLPGRRGNDGRPGLSSIKGQRGDPGFDGRTGAQGPPGDLTETGTYGPKGFPGLPGPPGQSGFPGPKGRPGDD
GISGYGQIKGLPGQPGQPGFAGAKGERGPSGSPGVSGSPAFIGAKGEKGSYGVRGFPGQTGRAGRPGLVSRKGSPGEIGF
PGPNGNPGNYGRKGERGTPGPPGRPGRYHPEVYLPGIPGDPGSSGRPGYSGPPGDLGFPGSPGLPGRPGNPGTPGREKGS

RGDPGTPVFPGLPGFPGPKGAPGINGFPGMKGEMGNRGASGPPGFHGESGQRGLIGEPGDQTGSPGFPGAKGERGDAGYP
GNAGPDGLPGNPGQDGIPGSNRTRGERGDAGVPGTHGPPGQSGLPGTPGGSGSEGLPGPPGPTASPGPPGSPGFPGSPGL
QGLNGLPGSKGSPGIPGFSLPGQKGEPGYPGPKGMLGDPGYGFGGPGFPGIKGTKGNSGVPGSRGFPGSPGPPGVLIGGY
PKGVVGDPGREGLPGPQGPSGPPGPPGQSTAFATKGDPGDPGINGGPGANGPPGDDGQAGTPGFPGQSGQKGGHGDQGLM
GFPGMKGHLGDNGYSGLKGDRGPTGDPGIRGPPGPVIAPGFVREPPPGPQGLKGAAGRIGAHGQRGSQGFIGPPGFKGGP
GRSGEPGRFGRPGNKGPRGDSGLPGRSGSTGGGGSPGQDGAPGLPGMPGRGVSFGLLLVKHSQSQEVPRCPLNMPRLWDG
YSLFYVEGSETAHNQDLGLAGSCLPRFNTMPFVYCNLNEVCNYGSRNDKSYWLSTNAPIPMMPVAEEAIREYISRCTVCE
APSVAIAIHSQGTAIPPCPRGYRSLWIGYSFLMHTAAGGEGGGQSLSSPGSCLEDFRSTPFVECQGARGTCHYFTNKYSF
WLTAIDENRQFDPEPVPETLKAGQPRTRASRCQVCMKNL
>Col4aB_[Eptatretus burgeri]
MQRWCLPLALCGLLIGTDAKVCRHAGCKTSGSSGIKGNKGKSGLPGTTGPPGLQGFPGPEGPHGPLGEEGSSGPPGRDGS
KGNRGPAGSPGFPGSSGLPGLPGQDGPLGPVGIPGCNGTKGEKGLPGGAGGFGPRGHRGQPGLPGTKGDSAKITGVLLPL
NGDKGIPGTHGQKGSPGRIGQPGPAGPSGSSGARGNDGPPGPPGEKGNVGLQFYGPRGSKGEKGFRGPPGRPASLEELKR
SPVQYEETYKGPKGESGDRGMPGPRGQAGIPGMSEPGMTGAKGEPGPLGQRGKPGKHGRGGSAGFDGIQGAPGESGRKGV
VGLKGGKGEQGMPGPPAQYDFDYNELEFRGDIGFPGLQGNKGEIGLTGPSGIPGFPGPKGEPYQGPAGDPGLPGEMGQKG
NGGLPGHSQLGPFGAPGHPGLPGQPGPRGPPGPKGTLPKGEQREHHINIGLPGLPGDKGFSGFPGEVGIKGDKGEVCLQC
TFAQNATRAPTGYPGLVGEPGVPGTDGSPGGKGDHGFTGTPGAAGPPGRPGSPGSTGLKGLKGESGRVDVEVFWEGDKGD
SGLPGSQGNPGRDGQPGIDGIPGNPGPKGEPALEGTKGDIGFPGPLGPPGISGEKGGQGLPSYGPSGFPGEKGDRGSHGT
PGVIGLLGQKGESGGTIETAGPLGPHGVRGEPGNVGSPGSNGLPGNDGIPGRQGPKGNQGSSGPRILSRPGEKGDYGPPG
RDGSPGLPSTPGKAGLPGLSGIPGMKGEPGVGLPGLPGLHGPPGLDGVPGAKGESAIPGIPGLPGQPGIDGALGQKGSQG
FPGPSGVPGLPGQKGKGLPGPAGPRGTRGPPGSDGLSGNPGQKGDRGLPGLGLPGKQGLKGAHGFLGVHGQLGDPGRPGK
PGNDGLPGINGAKGSMGIMGTPGPNGPPGPTGVPGLMGIKGDIGFVGPAGPPGLMGPVGDKGNPGLPGTPGILHPSLLQK
GEKGDQGDNGPAGRPGPKGNPGLLGETGSSGKGGFPGSPGGKGIKGDPGPPASPGIRGNRGLKGAMGEMGLPGSSGEVGD
TGLPGFRAEPGRPGHRGQKGDKGTAALPLPGLPGSPGMKGEAGWPGKPGLQGPKGNTGSHGLDGMAGGVGKKGEAGLPGF
PGTLGRRGEKGSPGLSGRSGVVGSAGPKGHRGEPGKAGPLGIPGVRGRDGFPGPVGPPGESGLTVSGLPGLSGPPGPQGP
KGQSGVPGSPGPPGHLGVKGEPGGAGRHGADGPPGRPGLPGMALPGVKGDPGVVGQRGLPGQLGLIGESGFPGFPGAKGD
KGLPGVSGMPGILGAKGNIGLPGRGGPHGRSGADGVKGNMGRSGIPGAVGLKGDQGLSGLSGPPGLKGQKGPEGPPGKGG
LALLPRITKADRGPTGTPGIHGDQGPPGPQGPFGPPGLLGPPGPTGMVGFPGLDGQRGKKGNRGFLGAFGEKGLDGPPGP
DGEPGMTGPPGGASIPHGFLVTRHSQTMFIPICPHGTTKVYDGYSLLHVQGNERAHGQDLGTAGSCLRRFSTMPFLFCNI
NNVCNFASRNDYSYWLSTPQAMPMNMAPISAPELQPFISRCAVCEASAMVIAVHSQAVTIPPCPSGWMSLWIGYSFVMHT
SAGAEGSGQALASPGSCLEEFRASPFIECHGRGTCNYYTNSYSYWLATVEGSKMFKKPESETLKAGNLRNRISRCQVCQR
RT
>Col4aC_4[Eptatretus burgeri]
MNKSVAFLAALALSLSSALLRSDAATCHGCASGSNCDCSGVKGDRGDRGVPGIQGRSGMPGFPGPEGPPGSEGPKGDLGD
SGVHGQKGHRGSPGMPGFAGTPGLPGLPGQDGPPGPPGIPGCNGTKGASGIPGDSLGGRRGPPGIPGHPGQKGDPGDVFS
TSGGIKGEQGLPGLSGRPGQRGTPGVTGTQGPPGPTGRPGSPGLPGPPGPKGTMGGDRLRFHGPKGEKGQRGLLGRPGPP
GSIGEQLGTREQFDLQPGPPGQKGEPGFPGERGLTGDSGPPGYGRKGEKGEQGDLGKRGKPGKDGEPGPPGFDGPPGQLG
RPGVNGRPGITGLKGEPGIQGGPGRVISGTGTQQGRGVKGDRGFPGPQGPKGEPGFRGQIGLPGPPGGTEGETFRGLPGL
PGPRGPQGPPGDPGRDGTSFPGQKGRDGIRGRAGAPGLPGPPGEPAYVPGPDFGSRGPPGRPGPPGEQGYPGERGFKGTK
GEHCLHCTTNGGRPPGIPGRPGPPGPQGFPGNPGLSGTKGDRGSIGIHGGPGSSGPPGNPGPMGMRGQKGDPGDAEGGYT
VKGDRGDPGHPGIRGLPGLNGLPGSAGIPGLPGPKGDPAPYGGKGERGFPGDPGTPGLPGERGTDGLPGSGFPGPHGMKG
VPGVPGSRGPPGVSGYKGEPGPTLPDSGLPGRPGPPGPPGAHGPPGSSGSPGNPGLSGFPGSKGEPGRSGFTPSGPPGPK
GERGPMGTPGLSGSPGRPGNDGRPGQDGFPGPKGESGIGRPGARGLPGPSGPSGFPGAKGDAGRPGLPGSSGLPGTDGFP
GTKGDPGLPGRPGSIGPPGLPGRGTPGLQGPQGPPGPAGSSGFPGSPGGKGTPGIPGRGIQGPPGENGQPGFPGSPGLKG
GPGFEGRPGSSGIPGSPGMKGDGGSSGFPGSSGPPGPQGEPGAARPGIKGNIGPPGFQGAQGSPGIQGPPGLPAAGGLKG
EKGNPGFSGSPGYAGAKGEPGFPGQPGGPGRPGFDGPKGDPGPAGTPGFPGSKGDPGRSGQSGFLGEKGFPGPQGSVGAP
GESGPSKIVKGAIGPPGFPGNNGPPGPAGLSGPPGPAGPVGPPGPPGSSGGPGFDGSPGVKGDVGNPGLSGERGFPGPQG
PAGLPGRIGSPGTGSMPHGFMLTRHSQMTDVPSCPAGTSVLYDGYSLLYVQGNERAHGQDLGTAGSCLRRFSTMPFMFCN
INNVCNYASRNDYSYWLSTPQPMPMSMEPIRGRDIQPFISRCVVCETPAMVIAVHSQSIMLPACPAGWVSLWIGYSFVMH
TSAGAEGSGQALASPGSCLEEFRSSPFIECHGRGSCNYYANSYSYWLSTIEPSEMFQKPSAETLKAGDLRSRISRCQVCM
RQQ
